# Cryo-ET reveals nucleosome reorganisation in condensed mitotic chromosomes *in vivo*

**DOI:** 10.1101/178996

**Authors:** Shujun Cai, Chen Chen, Zhi Yang Tan, Yinyi Huang, Jian Shi, Lu Gan

**Affiliations:** Department of Biological Sciences and Centre for BioImaging Sciences, National University of Singapore, Singapore 117543; Temasek Life Sciences Laboratory, National University of Singapore, Singapore 117604

## Abstract

Chromosomes condense during mitosis in most eukaryotes. This transformation involves rearrangements at the nucleosome level and has consequences for transcription, but the details remain unclear. Here, we use cryo-electron tomography to determine the 3-D arrangement of nucleosomes and other large nuclear features in frozen-hydrated fission-yeast cells. Nucleosomes can form irregular clusters in both interphase and mitotic cells, but they are smaller than expected for Hi-C domains. The nucleosomes are co-mingled with two features: nucleosome-free pockets and megadalton-sized “megacomplexes”. Compared to interphase, the nucleosomes in mitotic chromosomes pack into slightly larger clusters. However, nearest-neighbor distance analysis reveals that mitotic nucleosome clusters have the same internal packing density as in interphase. Furthermore, mitotic chromosomes contain fewer megacomplexes. This uneven chromosome condensation helps explain a longstanding enigma of mitosis: most genes are repressed but a subset is upregulated.

## INTRODUCTION

Chromatin structure influences key nuclear activities such as transcription, DNA repair and replication (Dixon et al., 2016). The fundamental unit of chromatin is the nucleosome, which consists of ~147 bp of DNA wrapped around a histone octamer Luger, 1997]. In mammalian cells, 2 - 35 nucleosomes pack into irregular “clutches” (Ricci et al., 2015) and more than 500 nucleosomes (calculated from nucleosome spacing) are thought to associate as topologically associating domains (Dixon et al., 2012). Likewise, in the fission yeast *Schizosaccharomyces pombe*, some 300 - 7,000 nucleosomes are thought to associate as compact globular chromatin bodies called domains (Mizuguchi et al., 2014; Kakui et al., 2017; Tanizawa et al., 2017). These studies suggest that chromatin higher-order structure arises from physical interactions of large groups of nucleosomes.

In mitotic cells, chromosomes condense into discrete structures that can be resolved in a light microscope. The factors involved in condensation have been well characterized (Hirano, 2016) and a number of models have been proposed for the large-scale organisation of chromatin domains (Maeshima and Eltsov, 2008). However, the molecular details of chromatin reorganisation are still unknown. Knowledge of how chromatin condenses in 3-D at the single-nucleosome level is needed to explain the nearly global transcriptional repression that happens to the mitotic cells of most eukaryotes (Struhl, 1998). Likewise, a 3-D model of chromatin could also explain how a subset of genes escape this mitotic repression and get upregulated (Rustici et al.; Oliva et al.; Peng et al.).

Some insights on *in vivo* chromatin organisation were made possible by new methods, including chromatin-conformation capture (Hi-C), super-resolution microscopy, and traditional EM of cells stained with DNA-proximal osmium (Beliveau et al.; Pombo and Dillon, 2015; Ou et al.). However, the resultant models are limited because these methods rely on population-averaged data, or have low resolution, or perturb the sample due to the fixation, dehydration and staining. Cryo-EM is a label-free method to visualize cells in a life-like frozen-hydrated state (McDowall et al., 1986; Eltsov et al., 2008). Cryo-electron tomography (cryo-ET) goes further and can reveal the 3-D positions of these complexes at ~ 4-nm resolution (Gan and Jensen, 2012). Using cryo-ET, we previously showed that nucleosomes in picoplankton and budding yeast pack irregularly, but we did not find evidence for mitotic chromosome compaction (Gan et al., 2013; Chen et al., 2016).

Fission-yeast mitotic-chromosome condensation resembles human chromosome condensation in both morphological and Hi-C phenotypes (Hiraoka et al., 1984; Naumova et al., 2013; Yam et al., 2013; Kakui et al., 2017; Tanizawa et al., 2017). We have now used cryo-ET to directly visualize chromatin both before and after mitotic condensation *in vivo* in the fission yeasts *S. pombe* and *S. japonicus*. To obtain cryotomograms with sufficient contrast to resolve nucleosomes *in vivo*, we imaged cells that were thinned by cryomicrotomy. To better understand the heterogeneous complexes in the crowded nucleoplasm, we also took advantage of recent advances in phase-contrast hardware and image-classification software (Bharat and Scheres, 2016; Khoshouei et al., 2017). We found that in interphase, nucleosomes frequently associate as small clusters or short chains. Nucleosome-free pockets and megadalton-sized “megacomplexes” are interspersed among these nucleosome clusters. In mitosis, chromosomes condense unevenly and have slightly larger nucleosome clusters. Mitotic chromosomes also have nucleosome-free pockets, but fewer megacomplexes. These phenotypes are conserved in both types of fission yeasts and lead to a model that explains how some genes can be transcriptionally upregulated in mitosis.

## RESULTS

### S. pombe subcellular structures are revealed at molecular resolution by cryo-ET

To understand how native chromatin organisation differs between interphase and mitosis, we imaged *S. pombe* cells by cryo-ET of frozen-hydrated sections (cryosections, ~ 90 - 130 nm nominal thickness). To ensure that the imaged cells had a known cell-cycle state, we arrested temperature-sensitive *cdc25-22* cells in G2 phase (Fantes, 1979) and cold-sensitive *nda3-KM311* cells in prometaphase (Hiraoka et al., 1984). Fluorescence microscopy confirmed that the latter cells have the well-known mitotic chromosome-condensation phenotype (Fig. 1A, S1). In a typical cryotomogram of an *S. pombe* cell, we could recognize organelles such as the endoplasmic reticulum, vesicles, and the nucleus due to their membrane morphologies (Fig. 1B and C). To further assess the quality of the cryosections and data, we performed subtomogram averaging of cytoplasmic ribosomes (Fig. S2A, B). The resulting average was similar to a low-pass-filtered budding-yeast ribosome crystal structure (Fig. S1C) (Ben-Shem et al., 2011). Therefore, the conformation of large complexes are preserved at the molecular level.

**Figure 1.**
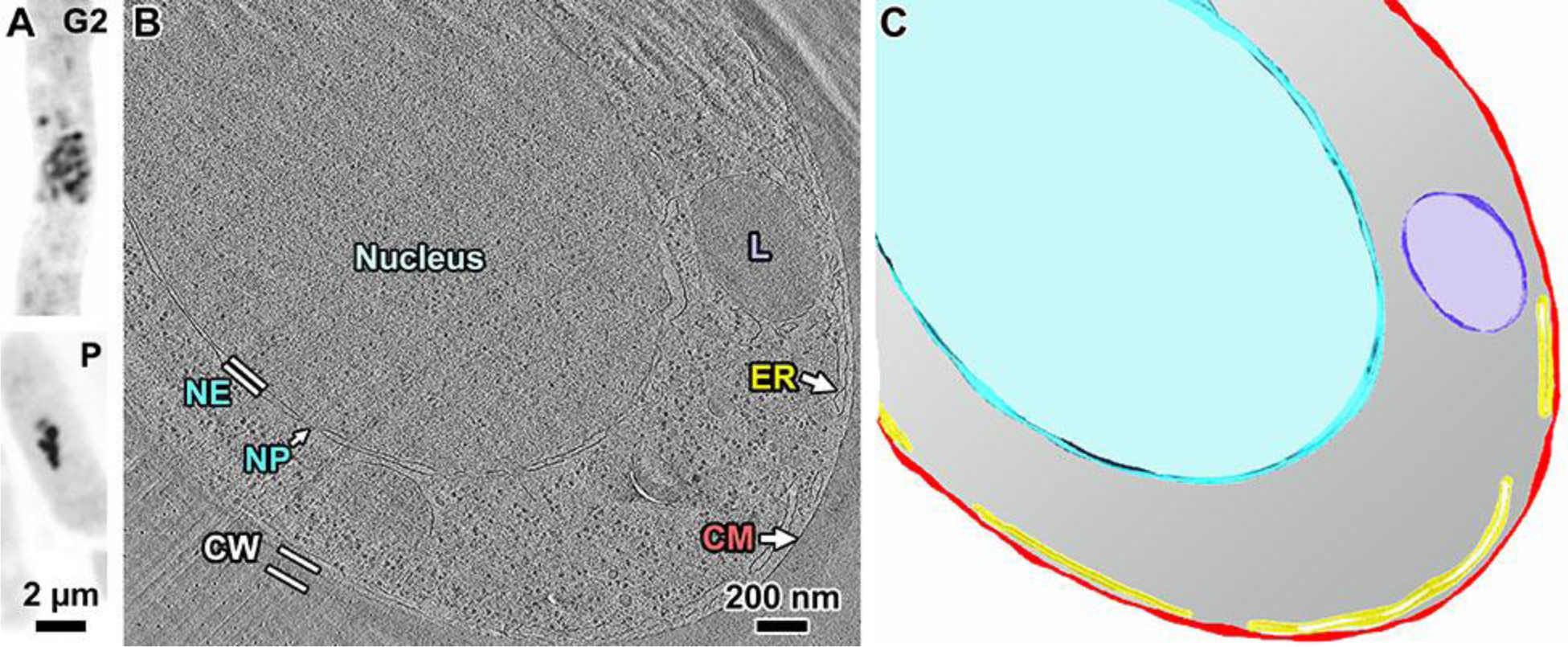
Strategy to study chromosome condensation in *S. pombe*. (A) Fluorescence images of DAPI-stained G2-phase and prometaphase (P) cells, with contrast inverted for clarity. (B) Cryotomographic slice (25 nm) of a G2-phase cell. NE: nuclear envelope; NP: nuclear pore; L: lipid body; ER: endoplasmic reticulum; CM: cell membrane; CW: cell wall. The wavy features in the upper portion of the cryotomogram are crevasses from sectioning; the thin parallel lines running from 2 to 8 O’clock are knife marks. (C) Segmentation of the cryotomographic slice in panel B, showing the cytological features with the same color scheme as the text labels from panel A.

### Interphase chromatin is arranged as loosely packed nucleosomes and irregular nucleosome clusters

G2-phase cell nuclei are filled with many nucleosome-like granular densities. For the sake of brevity, herein we call these densities nucleosomes due to their size, shape, abundance, location, and condensation phenotype all being consistent with nucleosomes (see below). Even if a minority of these densities are not nucleosomes, the conclusions we make below are still valid. Nucleosomes frequently associate as small irregular clusters less than 50-nm wide (Fig. 2B). Chain-like nucleosome configurations are also abundant (Fig. 2C). *S. pombe* linker DNA is only 7-bp, corresponding to ~ 2.5 nm (Lantermann et al., 2010), meaning that most nucleosomes should pack closely with at least two other nucleosomes. These two nucleosomes correspond to the −1 and +1 positions in the sequence. Because linker DNA is unresolved in our cryotomograms, we do not know if a cluster contains nucleosomes from a single or multiple sequences of nucleosomes. The remaining nucleosomes are not clustered together (Fig. 2D). Efforts to automatically identify nucleosome clusters with distinct motifs failed due to the irregular nucleosome packing. Small nucleosome-free “pockets” (< 50 nm) are also abundant (Fig. 2E). Densities much larger than nucleosomes (> 20 nm) are spread throughout the nucleus (Fig. 2F). These densities are consistent with being multi-megadalton nuclear assemblies such as pre-ribosomes, spliceosomes, and transcription preinitiation complexes (Gleizes et al., 2001; Allen and Taatjes, 2015; Oesterreich et al., 2016); we call these “megacomplexes” for brevity. Therefore, in interphase, nucleosomes associate as small irregular clusters that comingle with megacomplexes and nucleosome-free pockets.

**Figure 2.**
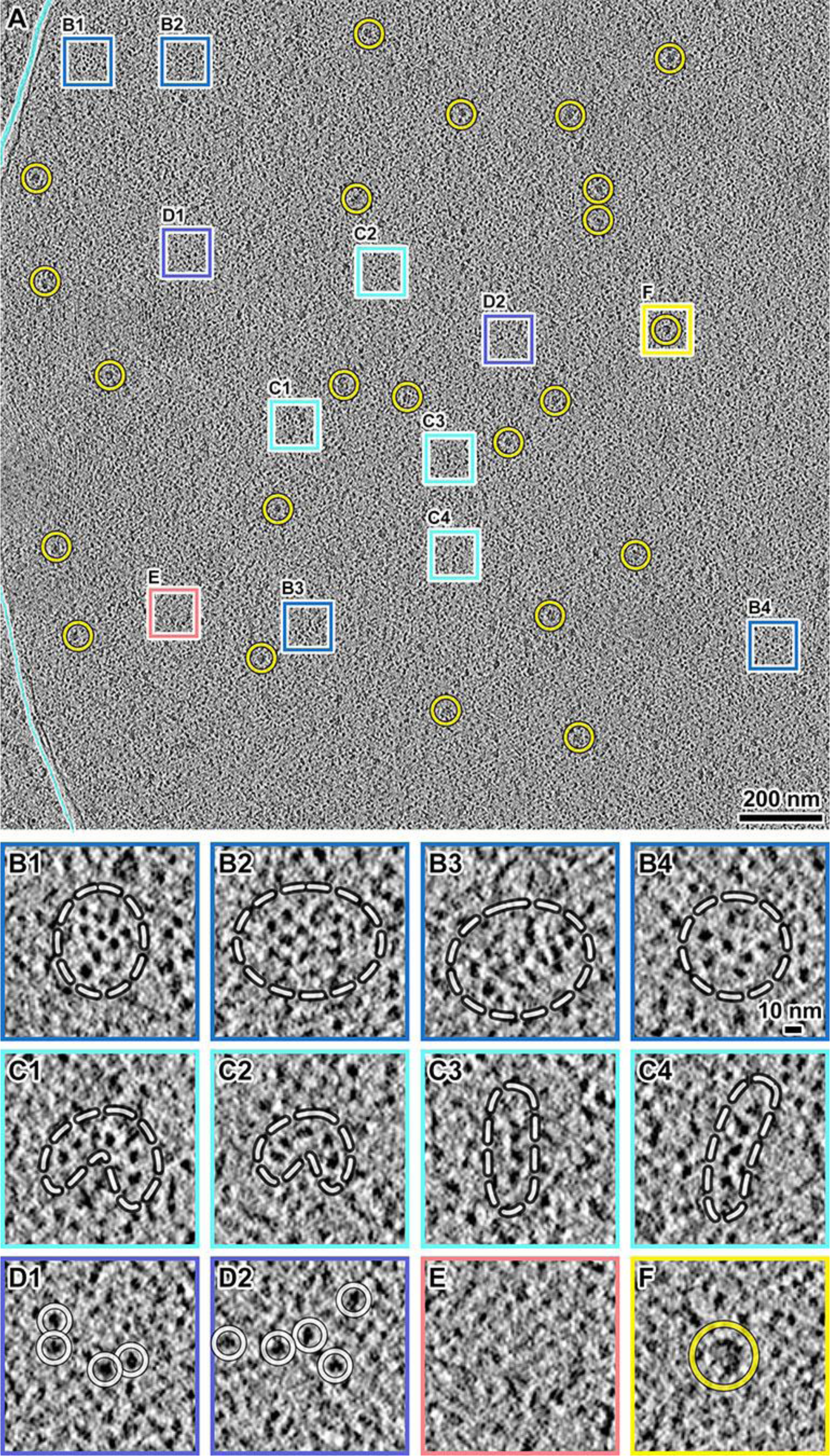
*S. pombe* G2-phase chromatin consists of irregularly packed nucleosomes and dispersed megacomplexes. (A) Cryotomographic slice (10 nm) of an interphase nucleus. Yellow circles: a subset of megacomplexes. The cyan lines denote the nuclear envelope. (B1 - B4) Four examples of nucleosome clusters. (C1 - C4) Four examples of nucleosome chains. In panels B and C, the white dashed lines delineate the approximate boundaries of the nucleosome clusters. Owing to the close packing and particulate nature of nucleosomes, it is not possible to annotate the exact “outline” of a nucleosome cluster. It is also not possible to determine how the nucleosomes are connected by linker DNAs, which generally cannot be seen. (D1 - D2) Two examples of loosely packed nucleosomes, which are circled. (E) An example position that has few nucleosomes. (F) An example megacomplex (circled).

### Condensed mitotic chromosomes contain fewer megacomplexes

Unlike in G2-phase cells, in each cryotomogram of a prometaphase cell we found a single position that has many features consistent with its being a section through a condensed chromosome. For example, nucleosome-like densities are abundant within these positions, but much rarer outside (Fig. 3A-C). Compared to G2-phase cells, in which abundant megacomplexes are interspersed with the chromatin (Fig. 2A), megacomplexes are mostly absent from these large contiguous regions in prometaphase cells (Fig. 3A). These positions are hundreds of nanometers wide, matching the size of condensed chromosomes seen by fluorescence microscopy (Fig. 1A, S1). If these positions are indeed chromosomes, there should be a large contiguous volume in the cell with few megacomplexes. To test this hypothesis, we sampled a larger nuclear volume by cryo-ET of 5 nearly sequential cryosections (summing to ~ 700 nm thick) of the same cell (Fig. S3A). We saw a single megacomplex-poor region in each of these cryosections, which is consistent with these positions being sections through one or two of the three fission-yeast chromosomes (Fig. S3B-F). Taken together, we located mitotic chromosomes and found they have fewer megacomplexes within.

**Figure 3.**
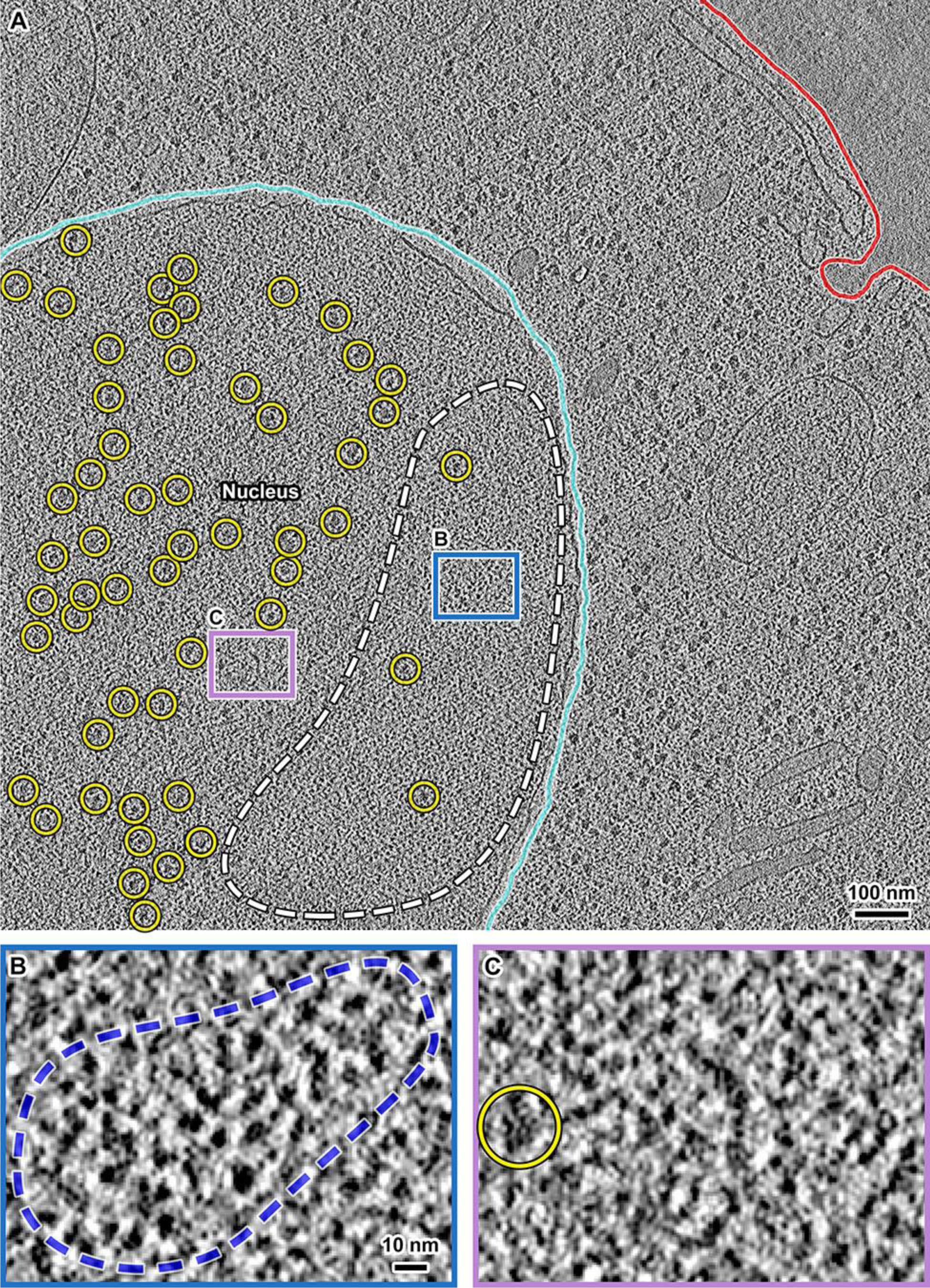
Condensed *S. pombe* mitotic chromosomes contain few megacomplexes. (A) Cryotomographic slice (11 nm) of a prometaphase cell. Cyan line: nuclear envelope, red line: plasma membrane, yellow circles: megacomplexes. The position circled with the dashed white line is a condensed chromosome. Rectangular boxes are enlarged 6‐ fold in panels B and C. (B) An example position with many nucleosome-like densities forming a cluster, circled with a dashed blue line. (C) An example position without nucleosome-like densities. Yellow circle: a megacomplex.

### Mitotic chromosomes condense unevenly and support transcription

In conventional (defocus) cryotomograms, we noticed that most nucleosomes appear more densely packed in prometaphase cells (Fig. S4). To better understand how chromatin is organised, we imaged cells by Volta phase contrast cryo-ET (Fig. S5). Volta cryo-EM data have much more contrast, making it possible to locate and determine the orientations of smaller protein complexes and to resolve protein complexes packed to near-crystalline density inside of cells (Engel et al., 2015; Khoshouei et al., 2017). The Volta cryotomograms confirmed what we learned from defocus cryo-ET: prometaphase nucleosomes more frequently packed into larger clusters (Figs. 4A and B, S5). Importantly, we confirmed the presence of some loosely packed nucleosomes and pockets in prometaphase cells (Figs. 4C and S5B, D), meaning that condensation is uneven. In summary, mitotic chromosomes consist of both densely and loosely packed nucleosomes.

**Figure 4:**
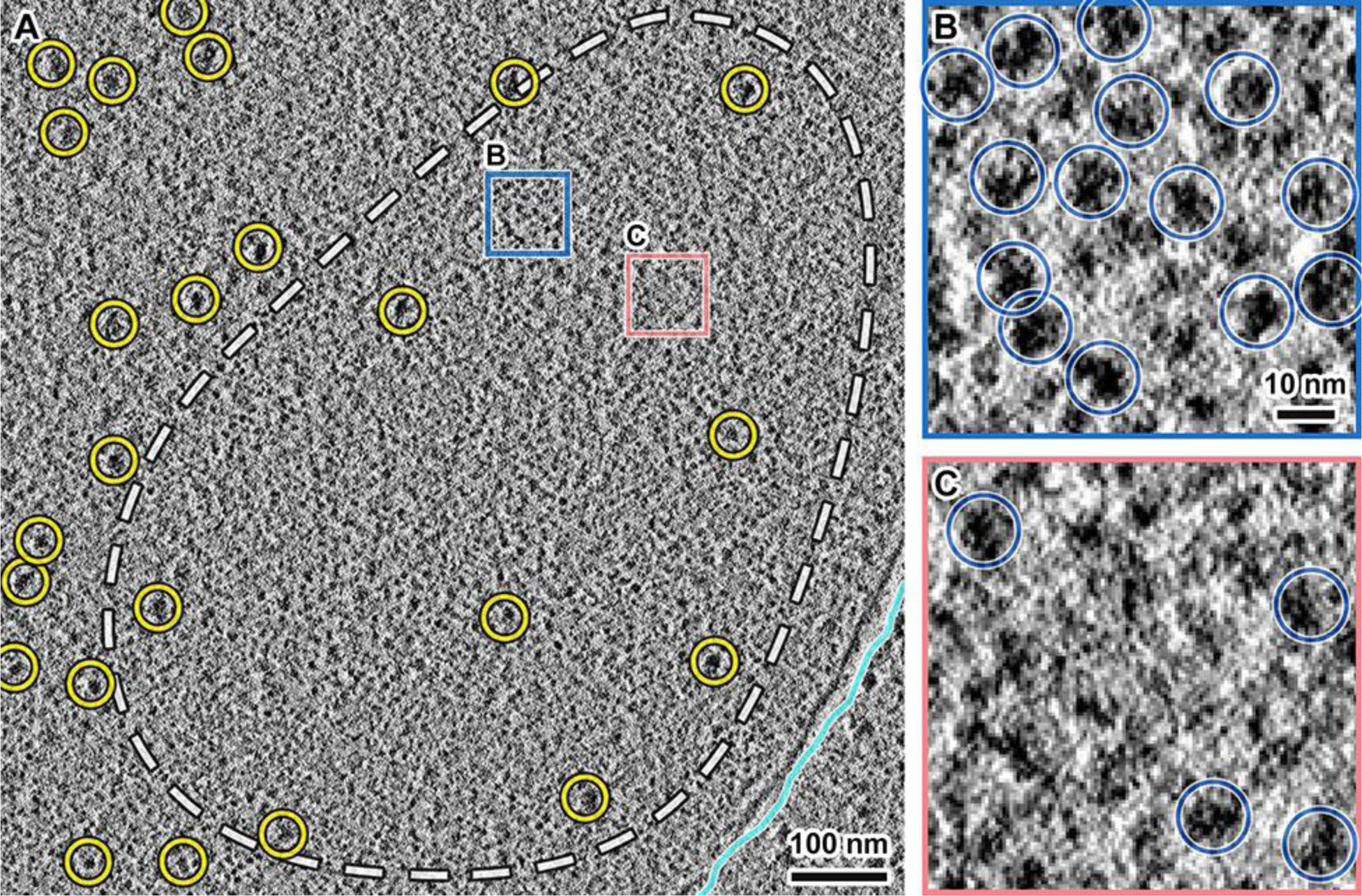
Larger nucleosome clusters and loosely packed nucleosomes coexist within *S. pombe* mitotic condensed chromosomes. (A) Cryotomographic slice (11 nm) of a nucleus in a prometaphase cell, imaged with Volta phase contrast. White dashed line: approximate chromosome boundary. Yellow circles: megacomplexes. The positions in the blue and salmon boxes are enlarged six‐ fold in panels B and C, respectively. (B) A position within the mitotic chromosome that contains many closely packed nucleosomes. (C) A position within the mitotic chromosome that contains fewer, loosely packed nucleosomes. Blue circles: nucleosomes.

The existence of loosely packed nucleosomes in prometaphase cells suggests that mitotic chromatin is permissive to transcriptional machinery, which is supported by the observation that some genes are up-regulated during mitosis (Rustici et al., 2004; Oliva et al., 2005; Peng et al., 2005). However, these earlier studies could not determine if *S. pombe* undergoes active mitotic RNA polymerase II transcription or if some stable mRNAs are left over from G2 phase. Phosphorylation of RNA polymerase II at Serine 2 of the carboxy-terminal domain (CTD) heptamer repeat is a conserved marker of transcription elongation (Komarnitsky et al., 2000; Harlen and Churchman, 2017). Using immunofluorescence, we confirmed the existence of RNA polymerase II with phospho-Ser2 CTD, and therefore active transcription, in prometaphase chromosomes (Fig. S6). To rule out the possibility of abnormal transcriptional activity in the mutants, we imaged asynchronous wild-type cultures, in which a small minority of cells was undergoing mitosis. We also detected phospho-Ser2 CTD signal in both early and late mitotic cells. Therefore, transcription is not shut-down globally in mitotic *S. pombe* cells.

To analyse the nucleosome rearrangement more objectively, we performed template matching to find their 3-D positions in Volta cryotomograms. This analysis showed that nucleosomes formed clusters in both G2-phase and prometaphase cells, but appear more crowded in the latter (Fig. 5A, B). If chromosomes condense via a uniform accretion of all nucleosomes, the average nucleosome nearest-neighbor distance (NND) distribution should shorten. However, the NND distributions of nucleosome hits in G2-phase and prometaphase cells were indistinguishable (two-tailed t-test, p > 0.05) (Fig. 5C). While mitotic chromatin might not condense by compaction of nearest-neighbor nucleosomes, they might instead condense from changes in interactions between groups of nucleosomes. We tested this hypothesis by creating histograms of 10^th^ NNDs and found that the population shifted to shorter distances in prometaphase cells (two-tailed t-test, p < 0.001) (Fig. 5D), which explains the more‐ crowded appearance of prometaphase nucleosomes. Taken together, our analyses reveal that chromosomes condense by the merging of small nucleosome clusters, resulting in a closer association between distant nucleosomes (Fig. 5E, F).

**Figure 5:**
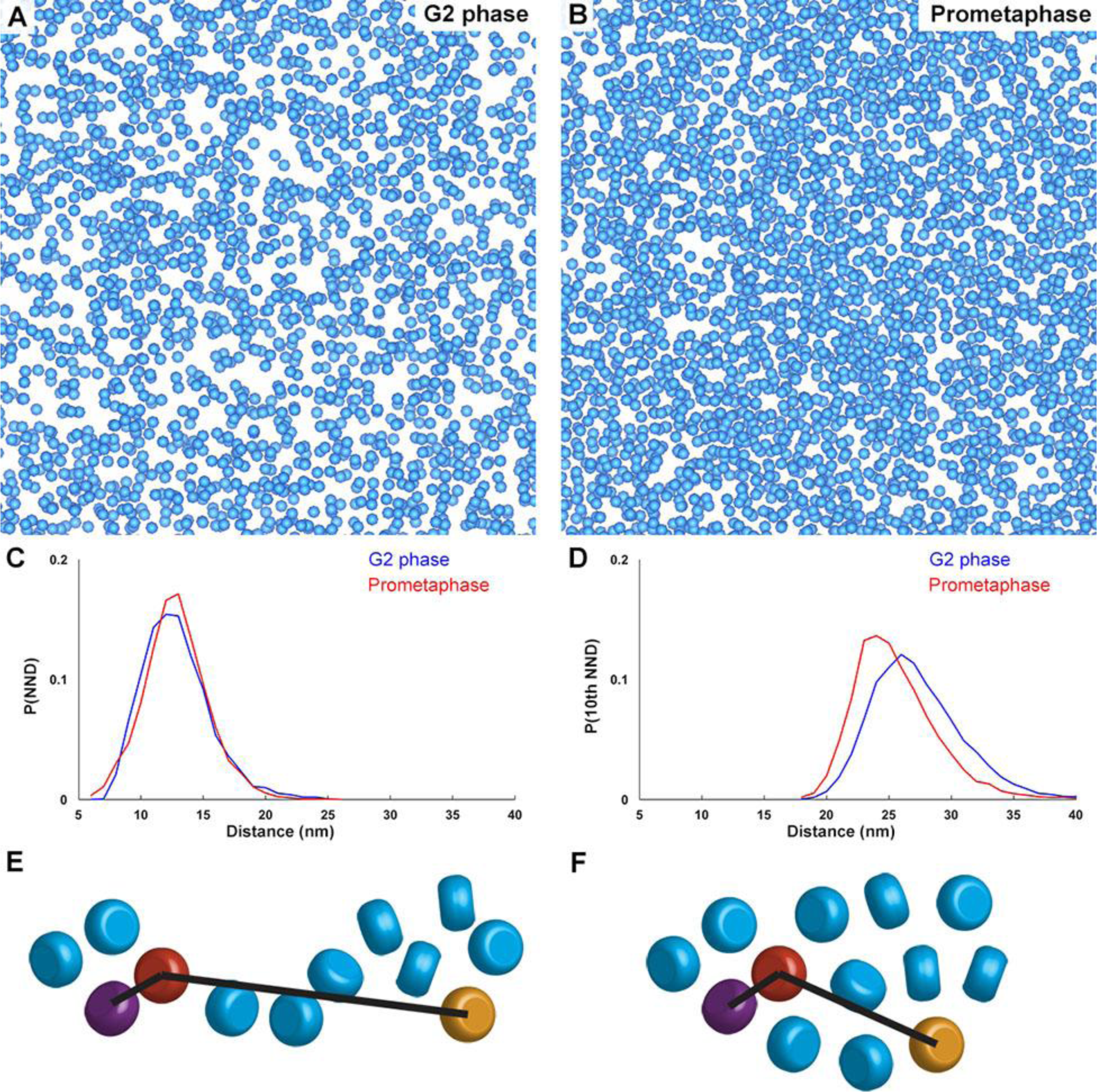
*S. pombe* nucleosome clusters are larger in prometaphase but have the same local packing density. (A and B) Template-matching hits of nucleosome positions (blue spheres) in Volta cryotomograms of a G2-phase (A) and a prometaphase (B) cell. (C) Nearest-neighbor distance (NND) analysis of template-matching hits. X-axis: NND, 1-nm bins, Y-axis: normalized probability. (D) Tenth NND analysis of template-matching hits. The X‐ and Y‐ axes are the same as in panel C. (E and F) Cartoons showing how a small (E) and a large cluster (F) of nucleosomes (rounded cylinders) could have the same NND, but different 10^th^ NND. In each panel, the purple and orange nucleosomes are the respective nearest and 10^th^-nearest neighbors of the red nucleosome. These cartoons do not show linker DNA, most of which cannot be seen in the Volta cryotomograms.

As an alternative approach to studying nucleosomes *in vivo*, we performed cryo‐ ET on the undigested chromatin from lysates of G2-phase and prometaphase *S. pombe* cells. Cryotomograms of cell lysis products can reveal more clearly the positions of nucleosomes (Cai et al., 2017). Subtomogram averages of nucleosome template‐ matching hits revealed unambiguous features of mononucleosomes including the groove between two DNA gyres (Fig. S7), demonstrating that our template-matching approach can locate most nucleosomes. In the lysates, there are many large nucleosome-free pockets wider than 50 nm in G2-phase chromatin. In contrast, prometaphase chromatin remained as more-compact masses with only a few, smaller pockets (Fig. S8). Therefore, prometaphase chromatin has denser nucleosome packing both *in vivo* and *in vitro*.

### *S. japonicusand* and *S. pombe* chromosomes condense by similar means

To test whether the formation of larger nucleosome clusters during condensation is conserved, we imaged *S. japonicus*, a bigger fission yeast that also undergoes mitotic condensation (Robinow and Hyams, 1989). *S. japonicus* has the same genome size (~ 12Mb) as *S. pombe*, but has a nucleus at least 8-fold more voluminous. Because *S. japonicus* is a less-developed model organism, we used asynchronous wild-type cells. These cells were frozen, cryosectioned, and subjected to Volta cryo-ET just like for *S. pombe*. To determine whether a cell was in mitosis or interphase at the time of freezing, we looked for cells with cytoplasmic microtubules, which exist only in interphase, or cells with nuclear microtubules, which exist only in mitosis (Yam et al., 2013). We found densely packed nucleosomes in *S. japonicus* mitotic chromosomes (Fig. S9A-D). Furthermore, NND and 10th-NND analyses showed that the nucleosome NND distributions were similar in interphase and mitosis (two-tailed t-test, p > 0.05), but that the 10th-NND distribution was shorter in mitosis (two-tailed t-test, p < 0.001) (Fig. S9E, F). Therefore, the formation of larger nucleosome clusters in mitosis is a conserved phenotype in fission yeasts.

### Dinucleosomes are irregular *in vivo*

The crowded nature of the nucleus makes it unfeasible to manually search for reproducible nucleosome-packing motifs. We therefore performed reference-free 2-D class-averaging analysis on di-nucleosomes extracted from our cryotomograms. Our previous 2-D classification of natural *S. cerevisiae* chromatin *in vitro* showed that abundant motifs can be found automatically this way (Cai et al., 2017). For example, pairs of nucleosomes sometimes packed face-to-face *in vitro*, which if present *in vivo* would limit access to regulatory positions like the acidic patch (Luger et al., 1997). However, the same analysis of di-nucleosomes in *S. pombe* did not reveal any classes containing face-to-face-packed nucleosomes (Fig. S10), meaning that such inter‐ nucleosome interactions must be extremely rare. Instead, di-nucleosomes were packed irregularly, including edge-to-edge packing, with ~ 3-nm gaps in between. G2-phase nucleosomes (Fig. S10A and C) were more irregularly packed in terms of the separation between candidate nucleosomes than in prometaphase cells (Fig. S10B and D). Taken together, our 2-D classification analyses did not find evidence of ordered nucleosome packing *in vivo*.

### A subset of nucleosomes may be partially unwrapped

In a rare instances, we were able to resolve densities that resemble linker DNA spanning two nucleosome-like densities in *S. pombe* G2-phase chromatin (Fig. S11A, B). Many of these linker-DNA-like densities were much longer than the average ~ 2 nm expected from MNase-digestion experiments (Fig. S11C-D) and nucleosome-mapping studies (Lantermann et al., 2010). Furthermore, in the NND analysis, there are a large number of nucleosome pairs that have NND values greater than 12 nm, i.e., the gaps between adjacent nucleosomes are wider than 2 nm (Fig. 5C). This inter-nucleosome gap was absent from tetranucleosomes reconstituted with the same linker DNA length as *S. pombe* (Ekundayo et al., 2017). One explanation is that the larger inter‐ nucleosome separation *in vivo* reflects the high end of the distribution of linker DNA lengths. Alternatively, some nucleosomes might be partially unwrapped – such as may be expected of fragile or pre-nucleosomes (Fei et al., 2015; Kubik et al., 2015), resulting in apparently longer linker DNA (Fig. S11E, F). The possibility of partially unwrapped nucleosomes led us to test whether the nucleosomes *in vivo* resembled mononucleosomes. To do this, we performed 2-D and 3-D classification of nucleosome template-matching hits (Bharat and Scheres, 2016). The 3-D classes taken from the Volta cryotomograms had the correct size and shape of nucleosomes, but they lacked the characteristic DNA gyres seen in the side view of the low-pass-filtered crystal structure of the mononucleosome (Fig. S12) (Luger et al., 1997). This difference might be due to either the insufficient signal-to-noise ratio of cryo-ET data or the conformational heterogeneity caused by biological factors such as partial unwrapping or both.

## DISCUSSION

We have directly visualized chromatin in both interphase and mitotic fission-yeast cells by cryo-ET. The condensation phenotypes are subtle. In both interphase and mitotic cells, most of the nucleosomes are packed into small irregular clusters. While the nucleosome clusters are slightly larger in mitosis, the nearest-neighbor distance does not shorten. In other words, there is no wholesale nucleosome accretion as commonly depicted in textbooks. The most notable ‐‐ yet still subtle ‐‐ changes are that mitotic chromatin contains fewer megacomplexes *in vivo* and that this chromatin remains more compact than interphase chromatin after being released from lysed cells. We propose a model of mitotic condensation that incorporates both structural and dynamic principles that takes into account recently published counter-intuitive Hi-C results (Fig. 6).

**Figure 6:**
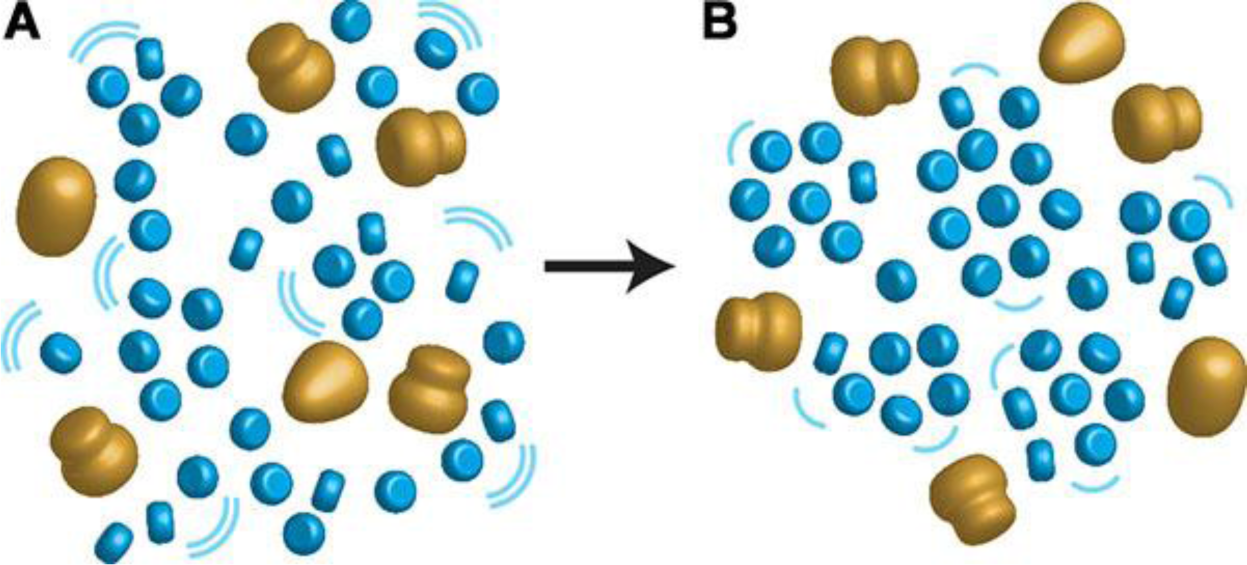
Mitotic chromatin is slightly more compact and less dynamic. (A) Interphase chromatin consists of both small nucleosome clusters and loosely packed nucleosomes. Blue pills: nucleosome; large gold bodies: megacomplexes. Megacomplexes (pre-ribosomes, spliceosomes, transcription preinitiation complexes) are interspersed between the nucleosomes and nucleosome-free pockets. (B) In condensed mitotic chromosomes, most nucleosomes form large clusters, thereby excluding or inhibiting the assembly of megacomplexes. Furthermore, chromatin dynamics (magnitude proportional to the size and number of the curved lines) also decrease. In both interphase and mitotic chromatin, ordered motifs such as face-to-face nucleosome-nucleosome interactions are largely absent.

### Cryo-ET reveals new insights into higher-order chromatin organisation

One of the long-standing goals of structural cell biology is to determine how a chain of sequential nucleosomes folds within minimally perturbed cells. Such a model requires the visualization of all the linker-DNA segments, but cryo-ET at present can resolve only a tiny minority of this DNA *in vivo*. Despite this limitation, the shortness of fission-yeast linker DNA (Fig. S10) tightly constrains our interpretation of our cryotomograms. Any given nucleosome density should have at least two adjacent nucleosome densities, which correspond to the -1 and +1 sequential positions. Each of these adjacent nucleosomes would be connected via short linker DNAs to another nucleosome, corresponding to -2 and +2 positions relative to the middle nucleosome. This line of reasoning fails for nucleosome-depleted positions, but these are in the minority. This linker-DNA imposed close packing supports our model that nucleosomes form small clusters and possibly small chains.

Our 2-D and 3-D classification analyses suggest that chromatin structure is irregular from the level of individual nucleosomes to mitotic chromosomes. Unlike the crystal structure (Luger et al., 1997), nucleosomes *in vivo* are largely conformationally heterogeneous. This heterogeneity may arise from interactions with small proteins, assembly/disassembly intermediates, partial unwrapping, or a combination of all these factors. Out of these possibilities, the partial unwrapping of *S. pombe* nucleosomes *in vivo* was predicted nearly a decade ago (Zlatanova et al., 2009). This variability in nucleosome structure and linker DNA length can then give rise to irregular oligonucleosome folding (Wong et al., 2007; Collepardo-Guevara and Schlick, 2014). Furthermore, the rarity of face-to-face nucleosome packing precludes chromatin-fiber interdigitation by nucleosome stacking as a major folding motif. Irregular chromatin is therefore a conserved feature in single-celled eukaryotes (Gan et al., 2013; Chen et al., 2016). A recent study using a new DNA negative-staining method concluded that chromatin structure is also irregular in mammalian cells (Ou et al., 2017), which is consistent with earlier EM studies of other mammalian cells (McDowall et al., 1986; Eltsov et al., 2008; Fussner et al., 2012; Nishino et al., 2012).

### Cryo-ET, Hi-C, and live-cell imaging paint a complex picture of chromatin

Hi-C is a “proximity-based” method wherein two DNA segments can be detected if they are close enough to chemically cross-link (Lieberman-Aiden et al., 2009). In this interpretation, Hi-C indirectly probes chromatin structure, and is revealing new principles of chromatin organization in *S. pombe* (Mizuguchi et al., 2014; Kakui et al., 2017; Tanizawa et al., 2017). A common Hi-C structural feature is the domain (also called TADs), a contiguous sequence in which nucleosomes are more likely to interact with other nucleosomes within this sequence than outside. *S. pombe* domains range from 40kb to 1Mb long, corresponding to 300 - 7,000 sequential nucleosomes. If domains are monolithic globular bodies as universally depicted in the literature, they should be visible in our cryotomograms as large, densely packed nucleosomes clusters. Such a cluster would be 7 - 20 nucleosomes wide, assuming a simple cuboid shape; other globular shapes would be equally large. The nucleosome clusters we saw were generally much smaller. We propose that small nucleosome clusters, which we see at the single-cell level by cryo-ET, correspond to different loci in each cell. If these clusters belong to Hi‐ C domains, they would sum to the much-larger structure due the effects of population averaging. As the simplest example, we illustrate how four sequential nucleosomes could give rise to a Hi-C structure that is larger than what can be found in individual cells (Fig. S13). Cell-to-cell variation may also explain differences between single-cell and population-based Hi-C studies (Nagano et al., 2013; Flyamer et al., 2017; Nagano et al., 2017; Stevens et al., 2017).

Two recent studies unexpectedly reported that short-range Hi-C contacts are less probable in mitotic *S. pombe* cells than in interphase cells (Kakui et al., 2017; Tanizawa et al., 2017), echoing a previous study of mitotic human cells (Naumova et al., 2013). Interestingly, quiescent B cells, which have condensed chromosomes, have lower short‐ range Hi-C contact probabilities than active B cells, which have decondensed chromosomes (Kieffer-Kwon et al., 2017). If Hi-C detections arise solely from nucleosome-nucleosome proximity, then the nucleosomes should move further apart in mitosis; we observed the opposite. What else, then, could explain the lower short-range Hi-C contact frequency in mitotic cells? A recent study found that within compact (160 nm) mammalian chromatin domains, all nucleosome motion is correlated (Nozaki et al., 2017). We propose that short-range Hi-C contacts in *S. pombe* and some mammalian cells is also dependent on chromatin dynamics (Hihara et al., 2012; Kakui et al., 2017). These dynamics slow down in mitotic cells due to the increase in nucleosome cluster size, and lowers the frequency that crosslinkable groups can interact with each other.

### The role of uneven condensation in transcriptional regulation

The strong correlation between mitotic chromosome condensation and transcriptional repression is well established (Taylor, 1960; Prescott and Bender, 1962). Mechanistic evidence of mitotic repression came from the observation that mitotic condensation is coincident with transcription-factor displacement (Martinez-Balbas et al., 1995). Yet, even though transcription is largely downregulated in mitotic *S. pombe* cells, (Oliva et al.), many cell-cycle-regulated genes actually get expressed more (Rustici et al.; Oliva et al.; Peng et al.). In agreement, our immunofluorescence experiments detected elongating RNA polymerase II in mitotic *S. pombe* chromosomes. We propose that chromatin structure at multiple size scales permits access to regulatory sequences, somewhat independently of dynamics. At the mononucleosome level, many chromatin‐ regulatory complexes recognize and bind to the nucleosome's face (McGinty and Tan, 2015). Because face-to-face nucleosome stacking is exceptionally rare in mitotic cells, this critical surface remains available for interactions. At the level of the chromosome, fewer megacomplexes are interspersed within the densely packed nucleosomes. However, uneven condensation leaves nucleosome-free positions that could remain permissive to the occasional passage or assembly of megacomplexes such as spliceosomes and transcription preinitiation complexes. If uneven chromosome condensation is highly conserved, the mechanisms proposed here could explain the recent observations of chromatin accessibility and transcription in mitotic mammalian cells (Hsiung et al., 2015; Teves et al., 2016; Palozola et al., 2017).

## MATERIALS AND METHODS

### Cell culture

*S. pombe* cell culture (Forsburg, 2003), G2-phase arrest (Ducommun et al.) and prometaphase arrest (Hiraoka et al.) were performed as previously reported. *nda3‐ KM311* cells were grown overnight in yeast-extract supplemented (YES) medium (30 g/L glucose, 5 g/L yeast extract, 225 mg/L adenine, 225 mg/L histidine, 225 mg/L leucine, 225 mg/L uracil) at 30°C with shaking. When the optical density at 600 nm (OD^600^) reached ~ 0.2, the cultures were cooled by incubation in a 20°C shaker. After a 10-hr incubation at this lower temperature, more than 90% of cells were arrested in prometaphase. *cdc25-22* cells were grown in YES medium at 25°C with shaking overnight. When the OD^600^ reached ~ 0.2, the cultures were warmed to 36°C in a water bath and then transferred to a 36°C shaker. After 4 hr, the majority of cells were arrested at G2 phase. *S. japonicus* cells were grown in YES medium at 30°C with shaking overnight until mid-log phase (OD^600^ ~ 0.6) (Yam et al.).

### Fluorescence microscopy

Cells were stained with 4′,6-diamidino-2-phenylindole (DAPI) following a published protocol (Toda et al.). Cells (1 ml) were fixed with 2.5% formaldehyde for 60 min at the restrictive temperature for mutants. The cells were washed twice by centrifugation at 1500 *x g* for 1 min and resuspension with distilled water. They were pelleted at 1500 *x g* for 1 min and stained by resuspension in 50 μl PBS, pH 7.4, containing 1 μg/ml DAPI. Ten μl of the sample was then added to a glass slide and imaged using a PerkinElmer Ultraview Vox Spinning Disc confocal microscope (PerkinElmer, Waltham, MA). Images were recorded using a 100x oil-immersion objective.

### Immunofluorescence

Log-phase cells (10 ml) were fixed with 3.7% formaldehyde for 90 min at 30°C (or at the restrictive temperature for arrested mutants). Cells were then pelleted by centrifugation at 1500 *x g* for 5 min. Cells were then resuspended in 1 ml PEM buffer (0.1 M PIPES pH 6.95, 2 mM EGTA, 1 mM MgSO^4^). Cells were washed once with 1 ml PEM this way and then resuspended in 1 ml PEMS (1.2 M sorbitol in PEM). Next, cells were spheroplasted for 15 minutes by incubation with lysis-enzyme cocktail (Abcam, cat# ab206997), diluted 1:1000 in PEMS at 30°C. Cells were washed with PEMS. The pellet was then resuspended in PEMS with 1% Triton X-100 and incubated at 22°C for 5 min. Cells were then washed twice with PEM and incubated in PEMBAL (PEM, 1% BSA, 100 mM L-Lysine hydrochloride) at 22°C for 1 hour. Cells were then incubated with rabbit anti-RNA polymerase II CTD-repeat YSPTSPS (phospho S2) antibody in PEMBAL (1:1000 dilution) at 22°C for 1 hour. Cells were washed three times with 1 ml PEMBAL and then incubated with Alexa Fluor 488-coupled donkey anti-rabbit antibodies at 22°C for 1 hour. Cells were then washed three times with PEM and resuspended in 50 μl of 1 μg/ml DAPI.

### Self-pressurized freezing

Self-pressurized freezing was performed based on a previous method, with modifications (Yakovlev and Downing, 2011). *S. pombe* cells were grown in YES medium until the OD^600^ reached 0.2 - 0.6. Cells were pelleted by centrifuging at 1500 *x g* for 5 min. Concentrated dextran stock (40 kDa, 60% w/v, in YES) was added to the cell pellet as an extracellular cryoprotectant to a final concentration of 30%. The cells were then quick-spun to remove air bubbles and then loaded into a copper tube (0.45 / 0.3 mm outer / inner diameters) with a syringe-type filler device (Part 733-1, Engineering Office M. Wohlwend GmbH). The tube was sealed by crimping both ends with flat-jaw pliers. The sealed tube was held horizontally, ~ 3 cm above the liquid ethane cryogen surface, and then dropped in. The flattened ends of the tube were then removed with a tube-cutting device under liquid nitrogen.

### Vitreous sectioning

Vitreous sectioning was performed as previously described with modifications (Chen et al., 2016). A perforated or continuous-carbon grid was coated with 10-nm gold colloids as fiducials for cryotomographic alignment. Gold solution (5 μl at 5.7 x 10^12^ particles/ml) in 0.1 mg/ml BSA was applied to the grid and then air-dried. Frozen-hydrated cells were cut into a 70-nm-thick frozen-hydrated ribbon using a 35° diamond knife (Cryo35, Diatome, Nidau, Switzerland) in a Leica UC7/FC7 cryo-ultramicrotome (Leica Microsystems, Vienna, Austria) at -150°C. Once the ribbon was ~ 3 mm long, the colloidal-gold-coated EM grid was placed underneath the ribbon. To minimize occlusion by grid bars at high tilt during cryotomographic imaging, the grid as aligned so that the ribbon was in between and parallel with the grid bars. The ribbon was then attached to the grid by operating the Crion in “charge” mode for ~ 30 seconds. The grid was stored in liquid nitrogen until imaging.

### Cell lysis

Log-phase *S. pombe* cells were pelleted by centrifuging at 1500 *x g* for 5 min and then spheroplasted for 15 minutes by incubation with lysis-enzyme cocktail at either 30°C or at the restrictive temperature for mutants. The spheroplasts were pelleted at 1500 *x g* for 2 min at 22°C and then lysed in 20 μl lysis buffer (50 mM EDTA and a 1:1000 dilution of protease inhibitor cocktail, Abcam, cat# ab206997) on ice for 15 min.

### Plunge freezing

Plunge freezing was done using a Vitrobot Mk IV (Thermo, Waltham, MA) operated at 4°C with 80% humidity. *S. pombe* cell lysates (4 μl) were applied to a freshly glow discharged perforated-carbon grid. The grid was blotted once (blot force: 1, blot time: 3 seconds) with filter paper (Whatman #1001055) and then plunged into liquid ethane.

### Cryo-electron tomography

Tilt series were collected using FEI TOMO4 on a Titan Krios cryo-TEM (Thermo, Waltham, MA) operated at 300 KeV and equipped with a field-emission gun, a Volta phase-plate device, and a Falcon II direct-detection camera. Details of the imaging parameters are shown in Table S1. Image alignment, CTF compensation, low-pass filtering, and 3-D reconstruction were all done using the IMOD software package (Kremer et al., 1996; Mastronarde, 1997; Xiong et al., 2009). Note that Volta cryotomograms were not CTF compensated. Cryotomograms were visualized as tomographic slices with 3dmod and as isosurfaces with UCSF Chimera (Pettersen et al., 2004).

### Template matching

Template matching was done using PEET (Heumann, 2016). A manually-selected subtomogram containing a nucleosome-like particle served as a template. The influence of the surrounding nucleoplasmic densities were suppressed by the application of a spherical mask. A search was performed on a grid with a 10-nm spacing within the nucleus. Template-matching hits within 6 nm were considered as duplicates and the extra hit was automatically removed. The cross-correlation coefficient cutoff criteria were the same as in our previous report (Cai et al., 2017). Briefly, the average cross‐ correlation coefficients of all template-matching hits was used as an initial cutoff. This cutoff was then manually adjusted (if needed) to minimize the number of detectable false positives and false negatives. To further remove false positives from megacomplexes, another round of template matching was performed using a ribosome like density selected within the tomogram as a template. The nucleosome hits that were within 12.5 nm of these megacomplexes were then removed with a Matlab script (available on request; Mathworks, Natick, MA).

For dinucleosome template matching, a manually-selected subtomogram of a di‐ nucleosome served as a template. A cylindrical mask (15-nm diameter, 22-nm height), rotated to align with the long axis of the dinucleosome template, was used to suppress the background surrounding densities. The search grid with a 25-nm spacing was generated within the nucleus. Owing to the missing-wedge artifact, nucleosomes have higher resolution in the X-Y plane. Therefore, the angular search was limited only to the Z axis. Hits within 15 nm were considered as duplicates. The cross-correlation coefficient cutoff criteria were the same as described above for mononucleosomes. False positives from megacomplexes fell into their own classes in the subsequent 2-D classification run (see below) and were removed.

### Reference-free 2-D and 3-D classification

Subtomogram classification and 3-D averaging of nucleosomes were done in RELION 1.4 and RELION 2.0 (Scheres, 2012a, b; Bharat et al., 2015; Kimanius et al., 2016), following the workflow of our previous study (Cai et al., 2017). The template-matching hit coordinates were imported into RELION and then particles were extracted with a 15.6-nm box and masked with a 13.5-nm-diameter sphere. For 2-D classification, the number of classes was set to 50 and the resolution of was limited to 3 nm. All nucleosome-like classes were selected. For 3-D classification, the number of classes was set to between 10 and 20, and the resolution was limited to 2-3 nm. Two to three rounds of 3-D classification were performed. False-positive were manually removed in between rounds.

For di-nucleosome 2-D classification, the box size and the mask diameter were set to 30 nm and 26 nm, respectively. As a default setting, RELION produces projections of all densities within the boxes, including the nucleoplasmic densities above and below dinucleosomes. To include only the dinucleosome densities, a python script (available on request) was used to generate projections from a 17.5-nm thick tomographic slice. The resolution of the data was limited to 2 nm to suppress the effects of high-resolution noise. The number of classes was set to 100, but many of the classes were very similar and therefore merged.

### Nearest-neighbor distance analysis

Nearest-neighbor distance analysis of template-matching hits (Figs. 5 and S9) was performed as previously described (Cai et al., 2017). The coordinates of the nucleosome hits were imported into Matlab. NND and 10^th^ NND were calculated using the Matlab function nearestneighbour.m (custom script available on request).

### Statistical analysis

Two-tailed t-tests for NND values were performed in Excel using the TTEST function. The number of NND values analysed for G2-phase and prometaphase *S. pombe*, interphase and mitotic *S. japonicus* were 10292, 11835, 9397, 4474, respectively.

### Data sharing

A cryotomogram, corresponding to Fig. 4, was deposited in the EMDataBank as EMD‐ 6846. The tilt series for all cryotomograms presented in this manuscript were deposited in the Electron Microscopy Public Image Archive as EMPIAR-10125.

## Acknowledgements

We thank the CBIS microscopy staff for support and training. We thank Snezhka Oliferenko for advice on *S. japonicus* culture and manuscript feedback; Mohan Balasubramanian for feedback and sharing *S. pombe* strains; Kazuhiro Maeshima for feedback. *S. japonicus* strains were obtained from JapoNet. SC, CC, ZT and LG were supported by NUS startups R-154-000-515-133, R-154-000-524-651, and D-E12-303-154-217, R-154-000-558-133, and MOE T2 R-154-000-624-112.

## Contributions

S.C - experiments, project design, writing, CC - training, YH - training, ZYT - experiments, JS - training, LG - project design, writing.

**Figure S1:**
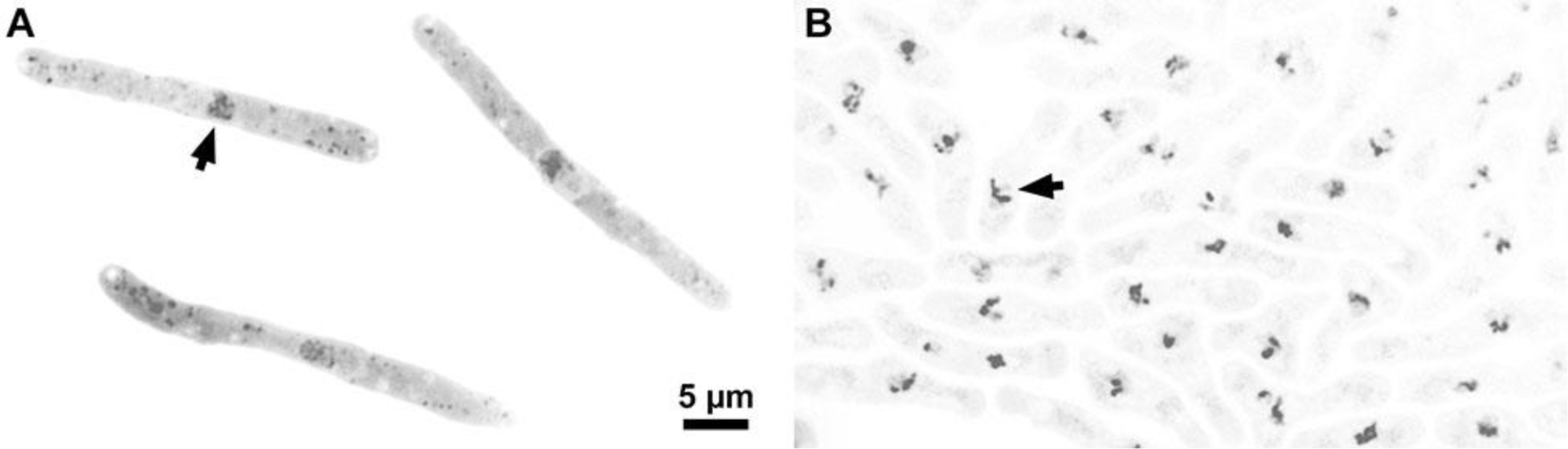
Synchronisation of *S. pombe*. (A and B) Fluorescence micrographs of DAPI-stained G2-phase (A) and prometaphase (B) cells. The contrast is inverted for clarity. Arrows point to an example nucleus in each panel.

**Figure S2.**
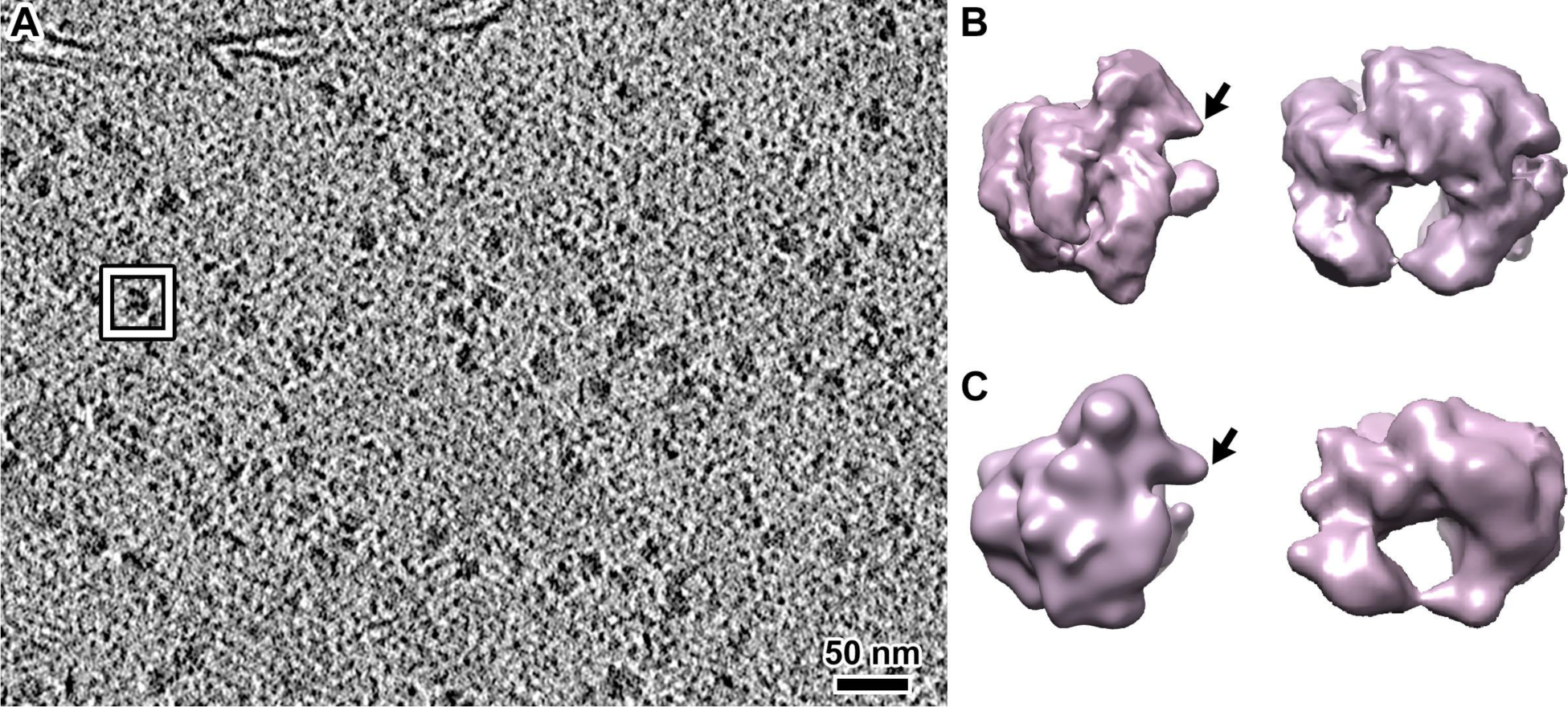
Cryotomograms reveal molecular detail *in vivo*. (A) Cryotomographic slice (25 nm) of a *S. pombe* cytoplasm position that contains many ribosomes, from the cell shown in Fig. 1B and C. (B) Subtomogram average of cytoplasmic ribosomes. (C) Crystal structure of *S. cerevisiae* 80S ribosome (PDB 4V88; (Ben-Shem et al., 2011)), low-pass filtered to 50 Ångstroms resolution. In panels B and C, the left image is related to the right image by a 90° rotation around the vertical axis, followed by a 180º rotation around the horizontal axis. Notice that in the orientation presented on the left, the “beak” motif (arrow) points to the right in both the simulated density map and the subtomogram average, indicating that the cryotomogram has the correct handedness. The two maps are not expected to be identical because the cytoplasmic ribosome map is an average of multiple conformational states.

**Figure S3.**
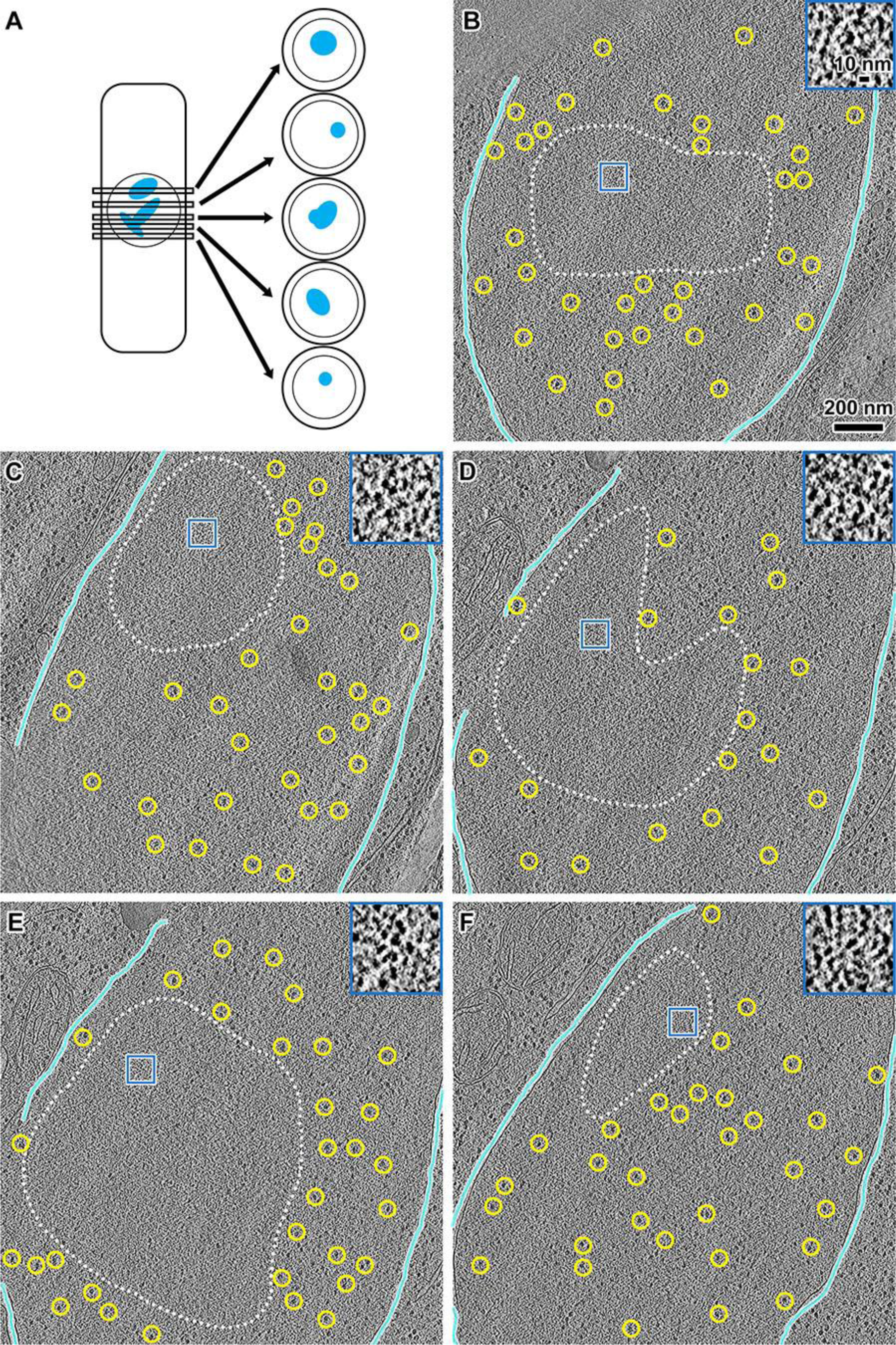
Serial cryotomography reveals the location of mitotic chromosomes. (A) A cartoon showing nearly serial cryosections of a cell. Some cryosections were not imaged due to ice contamination, damage, grid-bar blockage, etc. (B - F) Cryotomographic slices (10 nm) of nearly serial cryosections from a single prometaphase cell. Cyan lines delineate the nuclear envelope; there are gaps where the nuclear envelope densities were ambiguous. Dotted lines: approximate boundaries of condensed chromosomes. Yellow circles: examples of megacomplexes. For clarity, the few megacomplexes inside the dotted lines lines are not annotated. Inset: four-fold enlargements of blue boxes.

**Figure S4.**
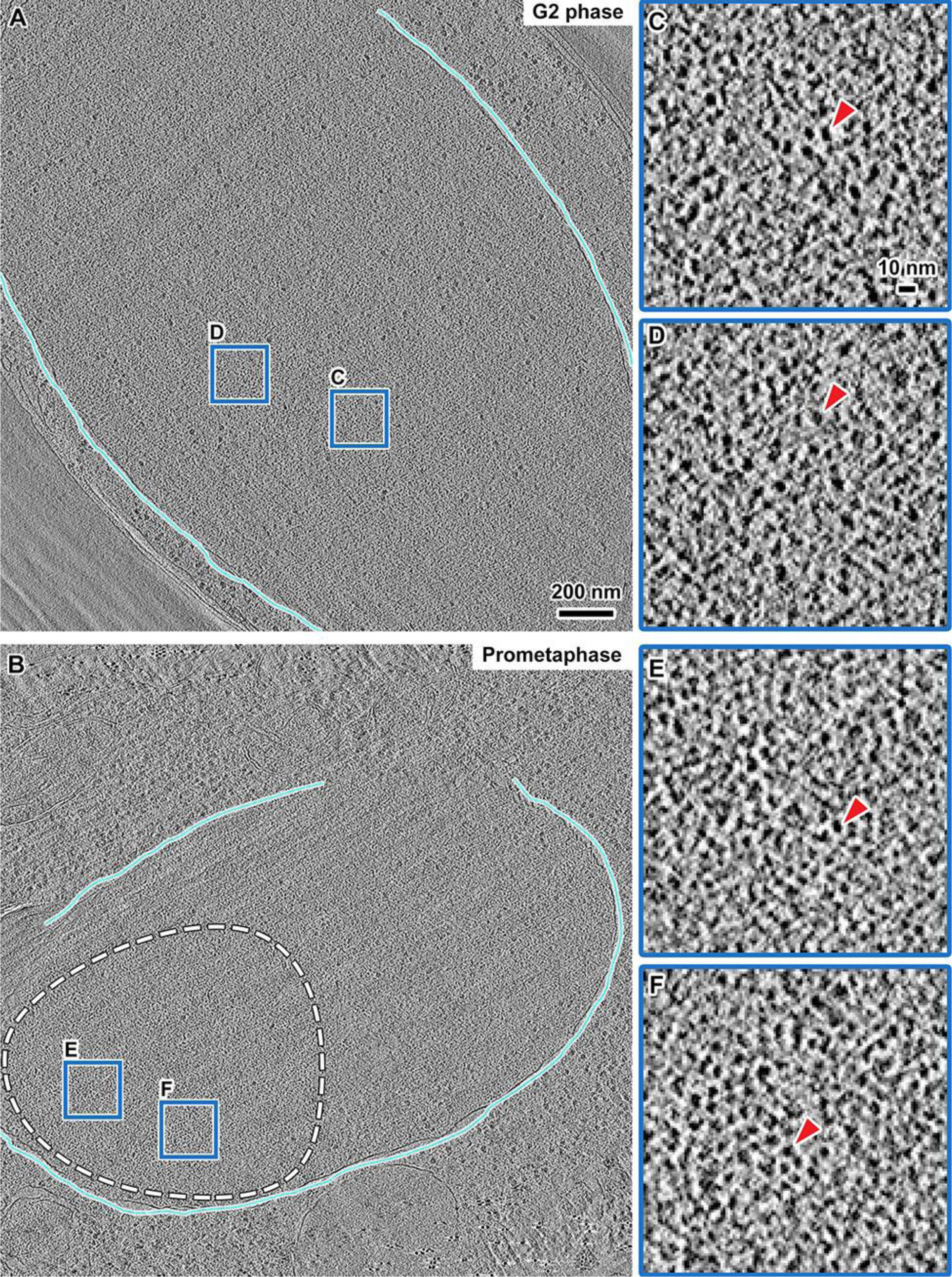
An example of nucleosome reorganisation in prometaphase cells. (A and B) Cryotomographic slices (11 nm) of nuclei in a G2-phase (A) and a prometaphase (B) cell. The cyan lines denote the nuclear envelope; there are gaps where the nuclear-envelope densities were ambiguous. Dashed circle in panel B: approximate outline of a condensed chromosome. (C - F) Six-fold enlargements of intranuclear positions that are enriched in nucleosomes, from the corresponding boxes in panels A and B. Red arrowheads point to nucleosomes.

**Figure S5.**
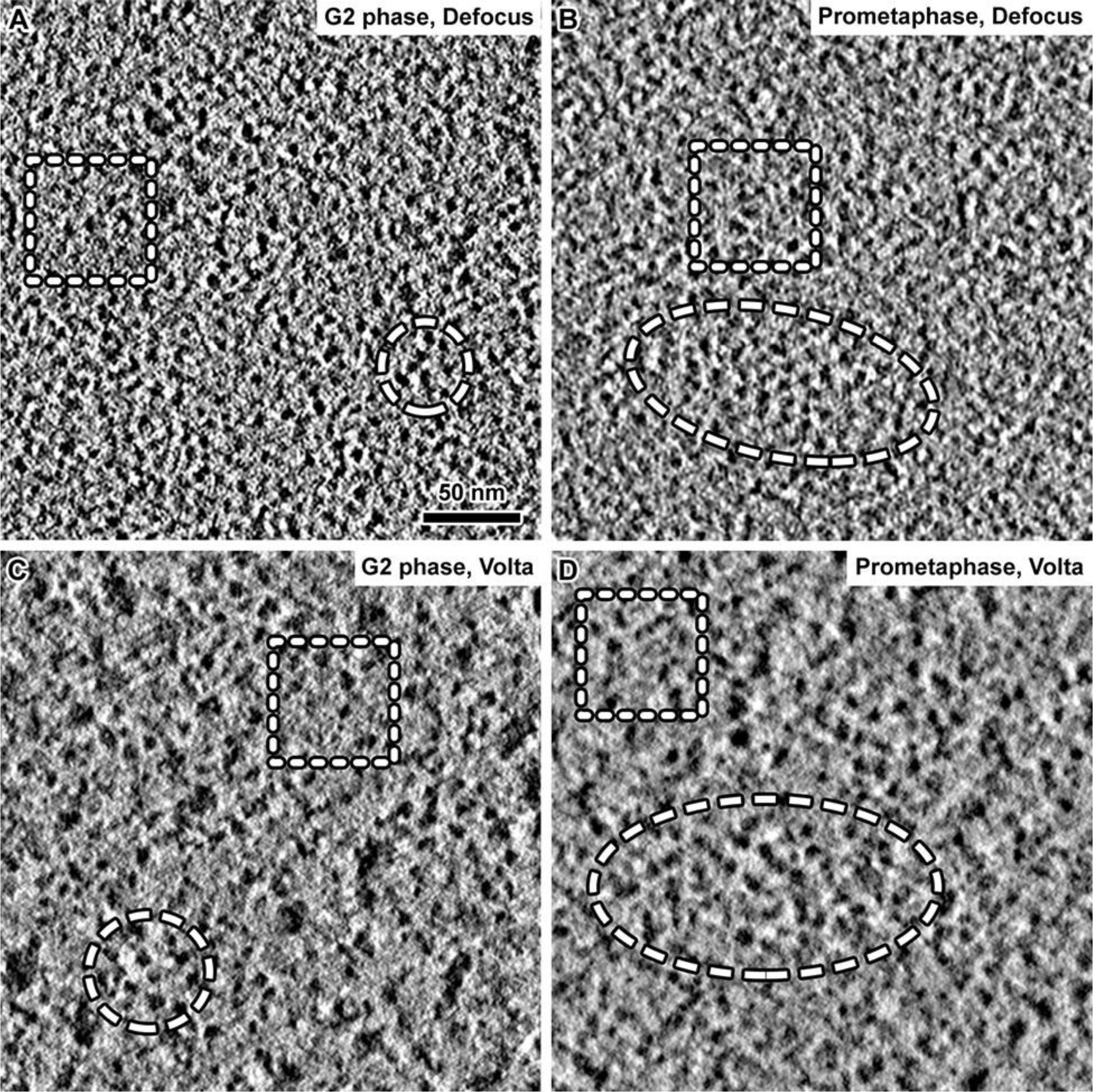
Nucleosome reorganisation is distinguishable in both defocus and Volta cryotomograms. (A and C) Cryotomographic slices (10 nm) of positions in G2-phase chromatin imaged with defocus (A) and Volta (C) phase contrast. (B and D) Cryotomographic slices (10 nm) of positions in prometaphase chromatin imaged with defocus (B) or Volta (D) phase contrast. Dashed circles: densely packed nucleosomes. Dashed squares: looselypacked nucleosomes.

**Figure S6.**
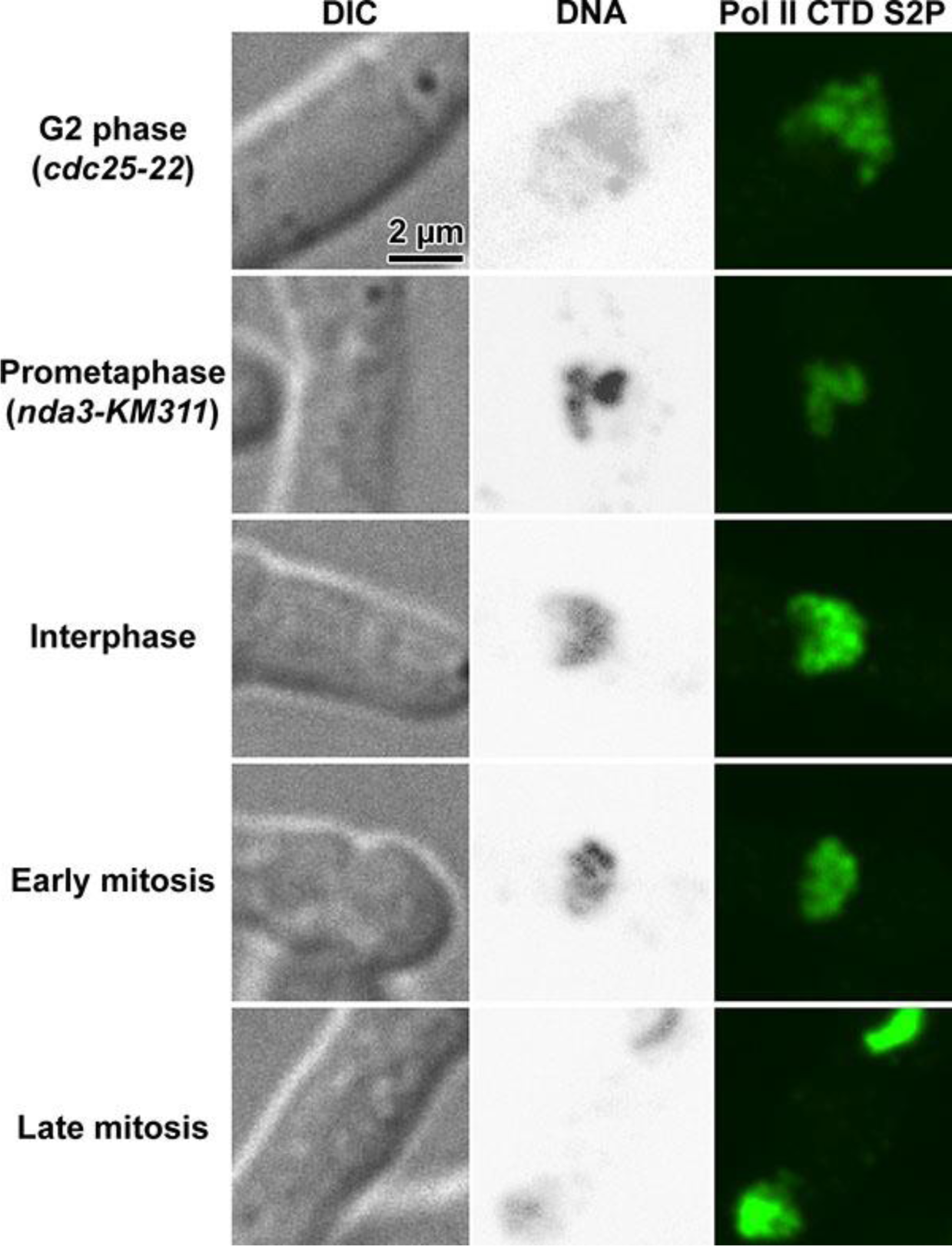
Transcription is not completely repressed during mitosis. DIC and fluorescence images of G2-phase, prometaphase and unsynchronized wild type cells. Left: DIC; Middle: DAPI-stained DNA with inverted contrast; Right: immunofluorescence of RNA polymerase II C-terminal domain repeat, phosphorylated at Serine 2 (Pol II CTD S2P). Top to bottom: *cdc25-22* cell arrested in G2 phase, *nda3‐ KM311* cell arrested in prometaphase, unsynchronized cell in interphase, unsynchronized cell in early mitosis, unsynchronized cell in late mitosis. In the unsynchronized cultures, the cell-cycle states were determined based on the chromosome morphology.

**Figure S7.**
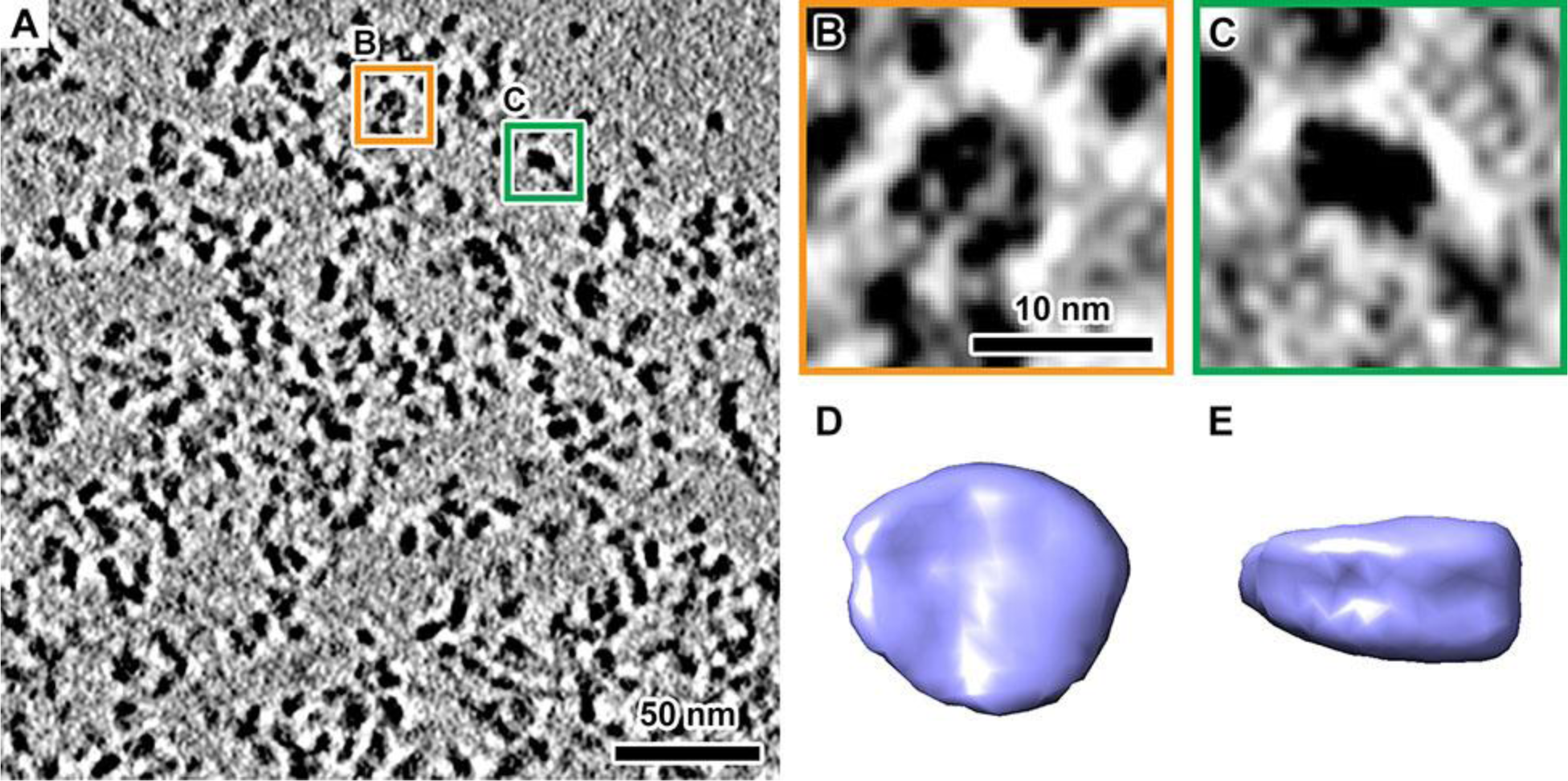
Nucleosomes remain clustered in cell lysates. (A) Cryotomographic slice (11 nm) showing the chromatin released from unsynchronised *S. pombe* cells. Nucleosomes oriented face-on (orange box) and side‐ on (green box) are enlarged 5-fold in panels B and C, respectively. (D and E): Face‐ and side-on views of a nucleosome subtomogram average of particles template‐ matched from the cryotomogram shown in panel A. Notice that DNA gyres are visible in the side view.

**Figure S8.**
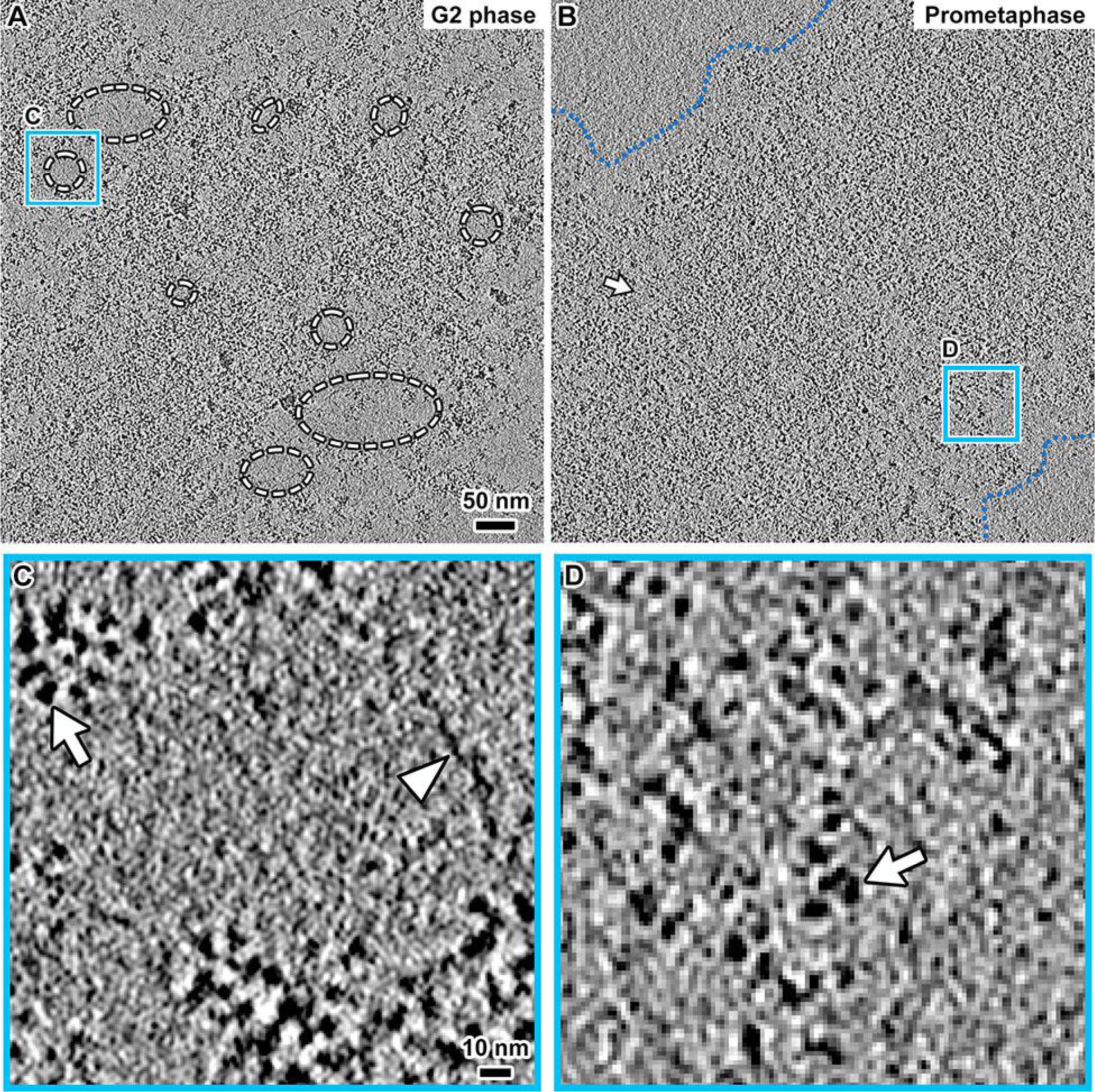
Figure S8. *S. pombe* prometaphase chromatin is also more compact than G2‐ phase chromatin *in vitro*. (A and B) Cryotomographic slices (30 nm) of chromatin from G2 phase (A) and prometaphase (B) cell lysates. Dashed circles in panel A: examples of large nucleosome-free gaps. Blue dotted lines in panel B: boundaries of chromatin mass. Arrow in panel B: examples of small nucleosome-free gap. (C and D) Four-fold enlargement of blue boxes in panels A and B, respectively. Arrows: nucleosomes. Arrowhead: DNA.

**Figure S9.**
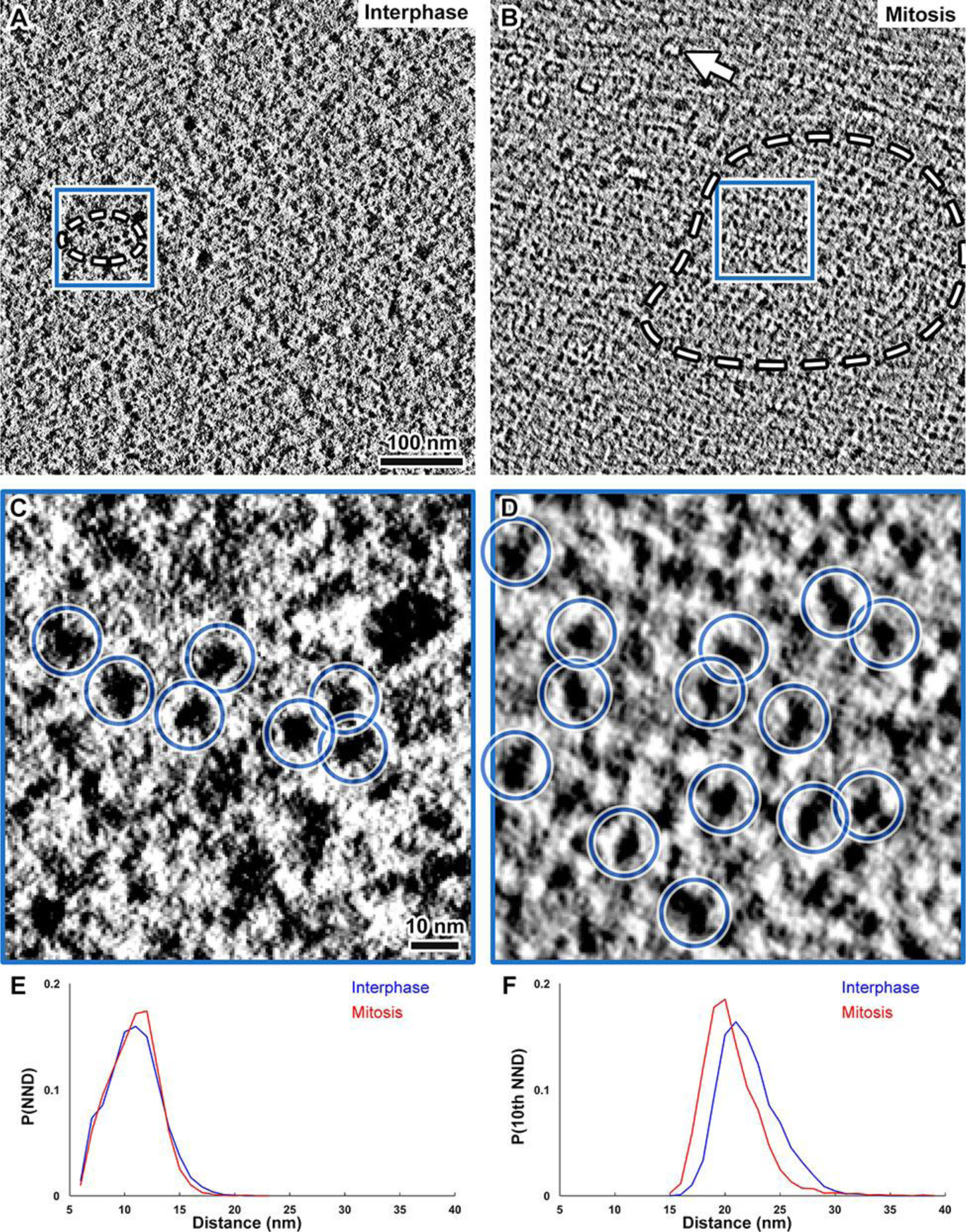
*S. japonicus* nucleosomes also pack into larger clusters in mitosis. (A and B) Volta cryotomographic slices (11 nm) of *S. japonicus* intranuclear positions in an interphase (A) and a mitotic (B) cell. Dashed circles: nucleosome clusters. Arrow in panel B: a spindle microtubule. (C and D) Five-fold enlargements of nuclear positions that have nucleosomes in panels A and B. Nucleosomes are circled in blue. (E) Nearest-neighbor and (F) tenth nearest-neighbor distance analyses of nucleosome template-matching hits.

**Figure S10.**
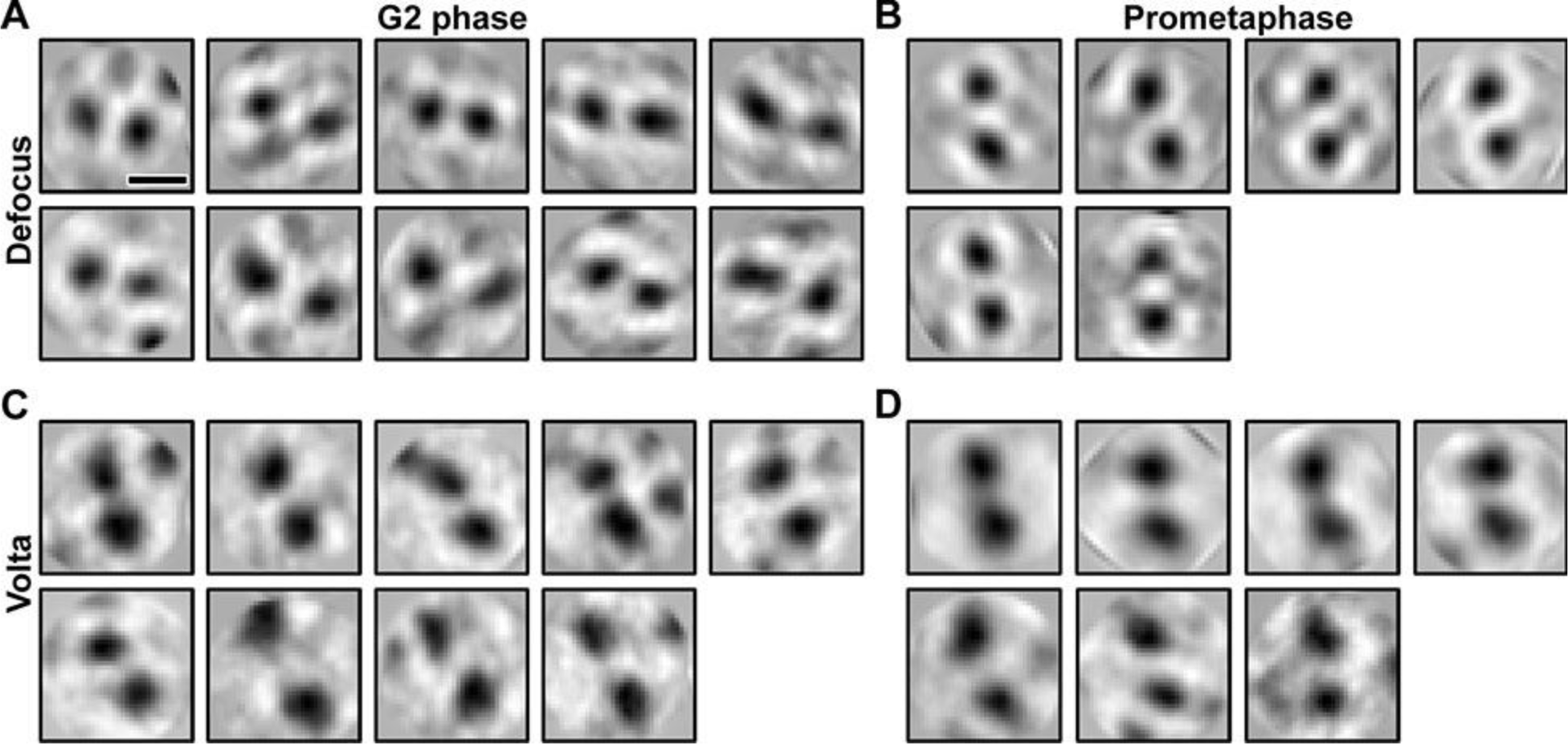
*S. pombe* nucleosomes do not stack face-to-face in either G2-phase or mitotic chromosomes. (A and C) Two-dimensional classification of dinucleosome template-matching hits in defocus (A) and Volta (C) cryotomograms of G2-phase cells. (B and D) Two dimensional classification of dinucleosome template-matching hits in defocus (B) and Volta (D) cryotomograms of prometaphase cells. Note that the 2-D classes showing megacomplexes (false positives) or tri-nucleosomes are not shown. Also note that some class averages, e.g., upper-left one in panel A, have a weak third density. This additional density may be from a lower-occupancy nucleosome position or a nucleosome that was truncated by the tomographic slice because it was not in the same exact plane as the other two. Scale bar in panel A: 10 nm.

**Figure S11.**
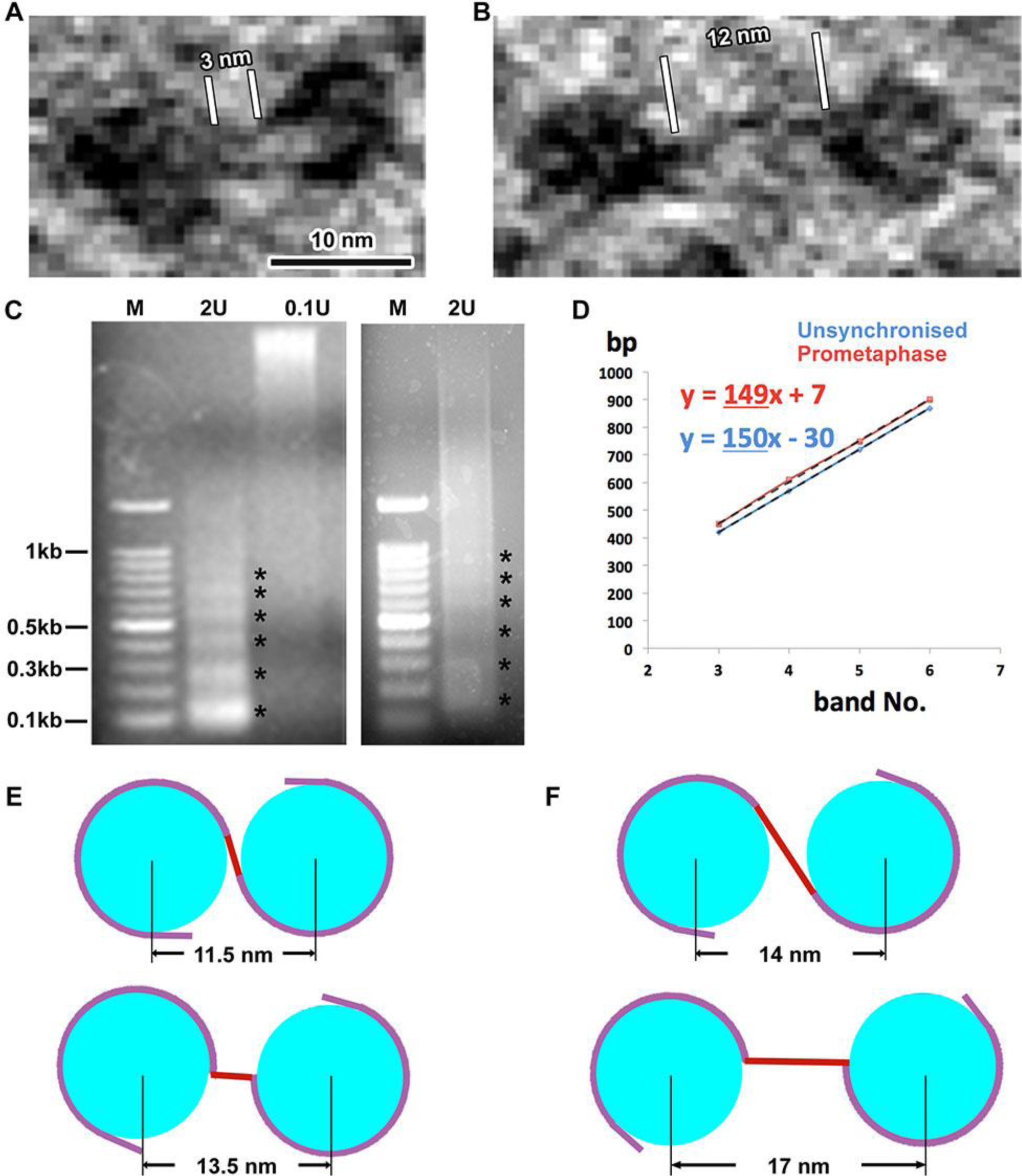
A subset of nucleosomes might be partially unwrapped *in vivo*. (A and B) Volta cryotomographic slices (10 nm) of a di-nucleosome-like density in a *S. pombe* G2-phase cell, showing densities consistent with short (A) and long (B) linker DNA. (C) The chromatin in unsynchronised (left; most cells are in G2-phase) and prometaphase (right) cells were fixed and then digested with 2 units or 0.1 units of MNase. DNA bands corresponding to mono‐ and oligonucleosomes are indicated by asterisks. (D) Nucleosome-repeat length plot of oligomer band number vs. base pairs. The monomer and dimer were excluded from the linear regression because the ‘nibbling’ of linker DNA at the ends would cause an overestimate of the slope and therefore, the nucleosome repeat length. (E and F) Cartoons showing fully (E) and partially (F) wrapped di-nucleosomes, drawn to scale for the nucleosome diameter and DNA length (but not DNA thickness). Red: *S. pombe* linker DNA. For the fully wrapped nucleosomes, the linker DNA is 2.5 nm (7 bp) long. For the partially wrapped example (10 bp unwrapped from the end of each nucleosome, the linker DNA becomes 6 nm long). The upper and lower panels show the extreme examples that minimize (upper) or maximize (lower) the inter-nucleosome separation for a given linker length. Upper panels: straight linker DNA. Lower panels: linker DNA bent at an energetically unfavorable ring angles at the entry/exit site.

**Figure S12.**
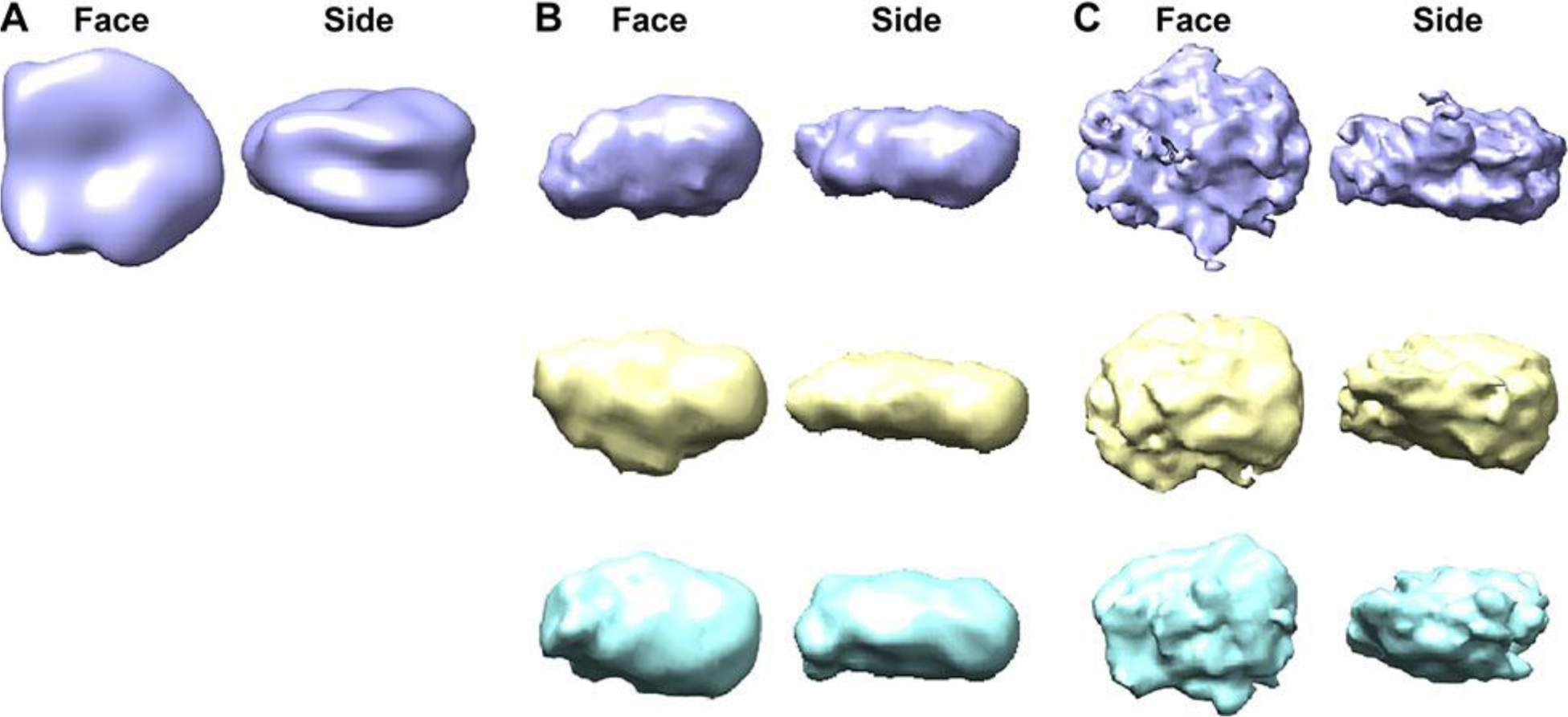
3-D class averages of nucleosome-like densities do not reveal high resolution mononucleosome-like features. (A) A simulated density map of a mononucleosome crystal structure (PDB 1ID3; (White et al., 2001)), low-pass filtered to 40 Ångstroms resolution. (B and C) Isosurface renderings of representative 3-D class averages of template-matching hits in the nuclei of a defocus (B) and a Volta (C) cryotomogram.

**Figure S13.**
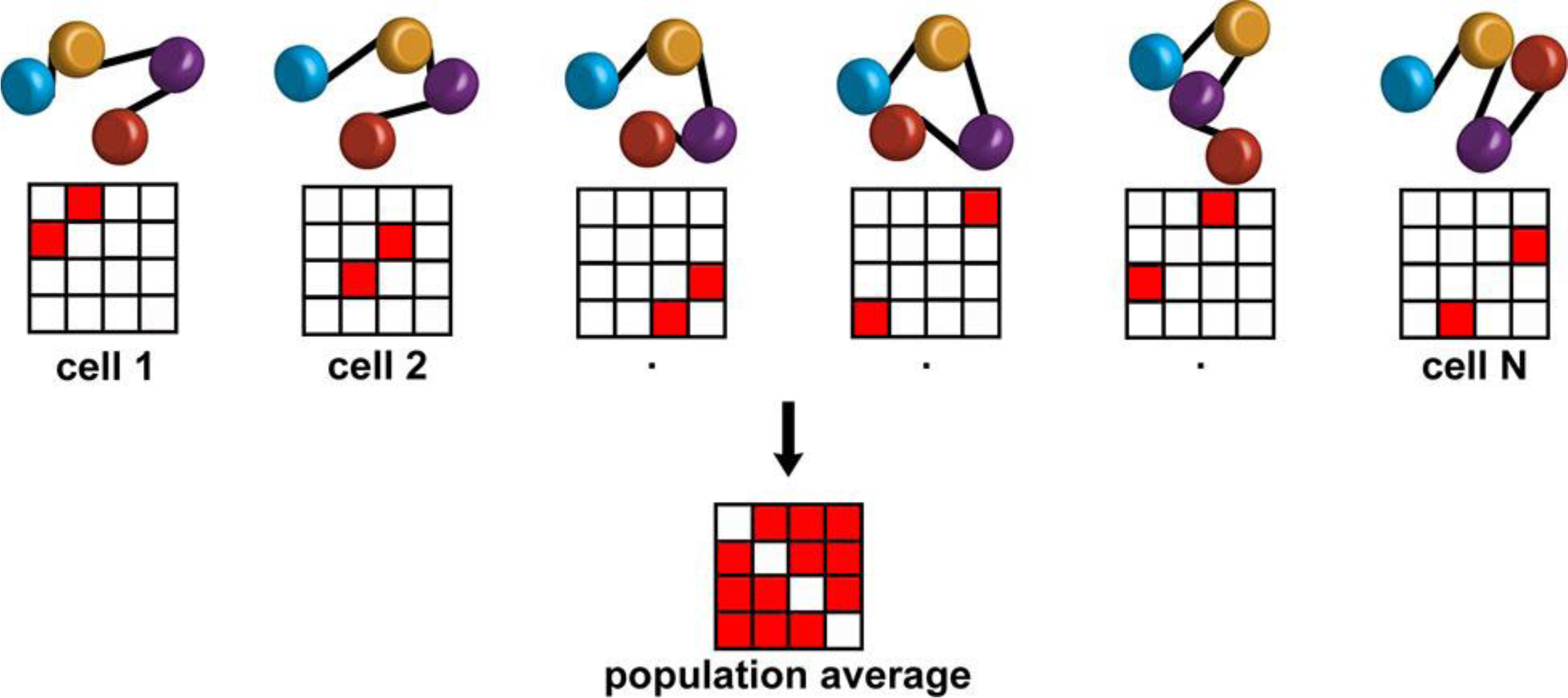
Reconciliation of *S. pombe* cryo-ET and Hi-C chromatin models. The blue, orange, purple and red rounded cylinders represent four sequential nucleosomes. Black lines indicate the connectivity between nucleosomes. Each column represents the nucleosomal arrangement of a different cell, at the time of fixation. The nucleosome pairs that are in physical contact have the potential to form crosslinks that can result in their detection, which is plotted in the contact matrices in the second row. Cell-to-cell variation at the time of fixation can result in the populational average shown in the Hi-C-based contact map (lower half). For clarity, only four nucleosomes are illustrated, but the concept here scales to clusters containing many more nucleosomes.

**Table S1.** Cryo-EM and image-analysis specifics.

|  |  |
| --- | --- |
| EM grid | C-flat 4-2-2, EMS continuous carbon |
| Microscope | FEI Titan Krios |
| Energy | 300 keV |
| Gun type | FEG |
| Camera | Falcon II Direct Detector |
| Tomography software | Leginon & FEI Tomo |
| Nominal magnification | 8,700, 11000, 18000 |
| Calibrated pixel size | 9 Å, 7.3 Å, 4.6 Å |
| Defocus (nominal) | -8-12 µm |
| Defocus (fitted) | -9-13 µm |
| Defocus (nominal, Volta contrast) | -0.5 µm |
| Cumulative dose | 80 - 150 e<sup>-</sup> / Å<sup>2</sup> |
| Dose fractionation | 1 / cosine |
| Tilt range | ±60° |
| Tilt increment | 1° or 2° |
| Tomogram processing & visualization | IMOD 4.9 |
| Template matching | PEET 1.10 - 1.11 |
| File format conversion | Bsoft 1.8.8 |
| Subtomogram classification & averaging | RELION 1.4, 2.0 |
| Density map visualization | UCSF Chimera 1.11 |
| Figure creation and compositing | Adobe Photoshop and Illustrator CS6, Google sheets |

**Table S2.** Cryotomogram details.

| Tomogram ID | Cells | Cell-cycle state, sample type | Figure | Dose | Pixel size (Å) | Defocus † (µm) |  | DPC/VPC |
| --- | --- | --- | --- | --- | --- | --- | --- | --- |
|  |  |  |  |  |  | nom | ref |  |
| 15Jun11_03 | *pombe* | G2, cell | 1B, C, S2 | 103 | 7.3 | 12 | 13-17 | DPC |
| 17May10_01 | *pombe* | G2, cell | 2, S5A, S10A | 80 | 7.3 | 12 | 13 | DPC |
| 17May09_18 | *pombe* | PM, cell | 3 | 80 | 7.3 | 12 | 12 | DPC |
| 15Nov16_10 | *pombe* | PM, cell | 4, 5B, D | 100 | 4.6 | 0.5 |  | VPC |
| 16Mar19_123 | *pombe* | G2, cell | 5A, C, S12 | 100 | 4.6 | 0.5 |  | VPC |
| 16Dec21_20 | *pombe* | PM, cell | S3B | 150 | 7.3 | 12 | 13 | DPC |
| 16Dec21_18 | *pombe* | PM, cell | S3C | 150 | 7.3 | 12 | 13 | DPC |
| 16Dec21_13 | *pombe* | PM, cell | S3D | 150 | 7.3 | 12 | 13 | DPC |
| 16Dec21_12 | *pombe* | PM, cell | S3E | 150 | 7.3 | 12 | 13 | DPC |
| 16Dec21_06 | *pombe* | PM, cell | S3F | 150 | 7.3 | 12 | 13 | DPC |
| 15Jun11_07 | *pombe* | G2, cell | S4A, C, D | 103 | 7.3 | 12 | 13-17 | DPC |
| 15May25_02 | *pombe* | PM, cell | S4B, E, F | 100 | 7.3 | 12 | 10-17 | DPC |
| 17May09_13 | *pombe* | PM, cell | S5B, S10B | 80 | 7.3 | 12 | 13 | DPC |
| 17May10_02 | *pombe* | G2, cell | S5C, S10C | 80 | 7.3 | 0.5 |  | VPC |
| 16Dec06_10 | *pombe* | PM, cell | S5D | 150 | 7.3 | 0.5 |  | VPC |
| 16Nov14_05 | *pombe* | Unsynchronised, lysate | S7 | 150 | 7.3 | 12 | 12 | DPC |
| 17Jul08_04 | *pombe* | G2, lysate | S8A, C | 108 | 7.3 | 12 | 14 | DPC |
| 17Jul08_01 | *pombe* | PM, lysate | S8B, D | 100 | 7.3 | 12 | 14 | DPC |
| 16Aug12_03 | *japo* | Interphase, cell | S9A, C | 100 | 5.8 | 0.5 |  | VPC |
| 16Jul25_14 | *japo* | Mitosis, cell | S9B, D | 100 | 7.3 | 0.5 |  | VPC |
| 16Dec06_7 | *pombe* | PM, cell | S10D | 150 | 7.3 | 0.5 |  | VPC |
| 15Nov16_09 | *pombe* | G2, cell | S11A, B | 100 | 4.6 | 0.5 |  | VPC |

Cell type: *pombe = S. pombe*; *japo = S. Japonicus*, G2 = G2 phase, PM = prometaphase, Dose, in electrons / Å^2^; † Nominal (nom) and refined (ref) defocus values; DPC = defocus phase contrast; VPC = Volta phase contrast

